# Highly Reversible Tunable Thermal-repressible Split-T7 RNA polymerases (Thermal-T7RNAPs) for dynamic gene regulation

**DOI:** 10.1101/2021.10.04.462087

**Authors:** Wai Kit David Chee, Jing Wui Yeoh, Viet Linh Dao, Chueh Loo Poh

## Abstract

Temperature is a physical cue that is easy to apply, allowing cellular behaviors to be controlled in a contactless and dynamic manner via heat-inducible/repressible systems. However, existing heat-repressible systems are limited and rely on thermal sensitive mRNA or transcription factors which function at low temperatures, lack tunability, suffer delays or overly-complex. To provide an alternative mode of thermal regulation, we developed a library of compact, reversible and tunable thermal-repressible split-T7 RNA polymerase systems (Thermal-T7RNAPs) which fuses temperature-sensitive domains of Tlpa protein with split-T7RNAP to enable direct thermal control of the T7RNAP activity between 30 – 42 °C. We generated a large mutant library with varying thermal performances via automated screening framework to extend temperature tunability. Lastly, using the mutants, novel thermal logic circuitry was implemented to regulate cell growth and achieve active thermal control of the cell proportions within co-cultures. Overall, this technology expands avenues for thermal control in biotechnology applications.

## INTRODUCTION

Temperature is a useful and unique physical cue that is non-invasive, highly penetrative, reversible, and easy to apply. Using these properties for gene regulation is useful in many engineering biology applications, such as in the bioproduction of proteins (Peng et al., 2012), the control of microbial therapeutics living *in vivo* (Piraner et al., 2017), and the development of microbial kill switches (Stirling et al., 2017). The use of temperature as a stimulus to regulate gene expression has advantages over the use of chemical which can be toxic to the cells (Aucoin et al., 2006; Briand et al., 2016), suffered from transport delay and lack of reversibility. With the increasing importance of the engineering of synthetic microbial community that is daunted by complex inter-population dynamics (Goers et al., 2014; Roell et al., 2019), the precise control of cell compositions with temperature can be a highly valuable tool to complement existing chemical based and optogenetic methods (Chen and Wegner, 2020; Guarino et al., 2020; Stephens et al., 2019). However, it is currently under-utilized due to lack of suitable temperature biosensors.

Microbial-based temperature biosensors can either be heat-inducible or heat-repressible, in which gene expression can be switched on or off at high temperatures respectively. While there are many developments in heat-inducible systems (Piraner *et al*., 2017; Rodrigues and Rodrigues, 2018; Valdez-Cruz et al., 2010), very few heat-repressible systems have been developed. Nonetheless, heat-repressible systems can be potentially useful in microbial-based bioproduction for phase-segregation of the biomass accumulation from induction, where the former occurred at higher temperatures to promote cell growth initially and in the latter phase, at lower temperatures to enhance enzyme stability and product yield in general (Peng *et al*., 2012; Qing et al., 2004; Singh et al., 2016; Sorensen and Mortensen, 2005). In addition, with the increasing need of temperature based genetic circuits to perform more complex functions in biotechnological applications, the use of multiple thermal sensors to form ‘thermal logic’ would be increasingly required (Piraner *et al*., 2017). This warrants further need for high-performance heat-repressible systems functioning in a temperature range (30 – 42 °C) suitable for microbial cell growth to work in tandem with the heat-inducible counterpart.

In general, reported microbial-based heat-repressible systems fall into two main categories, RNA-based and transcriptional-based regulations. In the first category, it relies on RNA thermometers which are temperature-sensitive regions on RNA molecules and with temperature shifts causing conformation changes to their secondary structures (Kortmann and Narberhaus, 2012). At its simplest, these structural changes of the RNA molecules, generally located in the 5’ untranslated region (UTR), can expose or occlude important sites such as the Shine-Dalgarno sequence and affect translation downstream. Several heat-repressible RNA thermometers which operate at different temperature ranges have been reported. These include examples such as the 5’ UTR of the cold shock protein A (CspA) mRNA (Keto-Timonen et al., 2016; Qing *et al*., 2004) which is stable at low permissive temperature (15 °C) to promote the translation of the gene of interest (GOI), and the RNAse E-cleavable RNA thermometer (Hoynes-O’Connor et al., 2015) which at the permissive temperature (27 °C) is hidden within a hairpin loop to prevent degradation of target mRNA. Generally, RNA thermometers can have almost immediate response to temperature shifts as they directly regulate the translation of existing or nascent mRNA strands (Kortmann and Narberhaus, 2012) and their structures have been increasingly predictable with the rendering from computation-guided designs (Hoynes-O’Connor *et al*., 2015; Sen et al., 2017). Nonetheless, they typically suffer from small fold inductions (< 10-fold) and broad thermal transition (> 5 °C) (Hoynes-O’Connor *et al*., 2015; Piraner *et al*., 2017; Sen *et al*., 2017; Yang Zheng et al., 2019) as brought about by possibly unstable equilibrium shifts between the open and closed RNA structures at different temperatures (Wei et al., 2016). The low activation temperature (15 °C) of some RNA thermometers while useful for improving protein solubility (Qing *et al*., 2004; Sorensen and Mortensen, 2005), may not be as suitable for bacterial growth and biocatalytic processes which required higher temperatures (> 30 °C).

In the second category, transcriptional regulation mainly relies on temperature-sensitive transcription factors that change their structural configurations and affect binding onto the operator or promoter sites that control gene expression downstream. There are several notable examples of transcriptional based heat-inducible systems as described (Fu et al., 2020; Hussaina et al., 2014; Pearce et al., 2017; Piraner *et al*., 2017). However, there are limited heat-repressible systems which rely on transcriptional regulation (Liang et al., 2007; Yang Zheng *et al*., 2019). A recent work has performed temperature-based screening on the modified bacteriophage λ Cl434 repressor that contained the tobacco etch virus (TEV) cleavage site (Yang Zheng *et al*., 2019). In the study, through mutagenesis, the CI434 repressor was converted from a heat-inducible to a heat-repressible form and achieved fold changes of 2 – 4 within 25 – 42 °C. To improve the expression at permissive low temperatures and to reduce leakiness at high temperature, a temperature-sensitive TEV protease that specifically cleaves the repressor at low temperatures and a mf-Lon protease that recognizes and degrades the output tagged reporter protein at the restrictive high temperature were introduced into the circuit. This significantly improved the performance of the overall system. The strategy of introducing proteases in a feedback configuration to actively degrade the repressors or target proteins to reduce leakiness could be useful to improve the performance of thermoswitches. However, because of the multiple modules involved, the system reportedly suffered from delays and would be difficult to achieve rapid ON and OFF dynamics useful for transient regulation.

Nonetheless, existing designs of heat-repressible thermoswitches are centred mainly on the use of heat-sensitive transcription factors or 5’ UTR of target mRNAs to regulate expression. The direct thermal control of the RNA polymerase that transcribes the GOI would offer an important alternative mode of thermal regulation in synthetic biology, with potential to create compact heat-repressible system, simplifying the implementation of synthetic circuits, and enable highly sensitive ON and OFF response to heat changes. The T7 RNA polymerase (T7RNAP) is a popular RNAP used in the field of biotechnology (Liu et al., 2018; Wang et al., 2018). The use of the phage polymerase is attractive because the polymerase can effectively decouple transcription from the host machinery and recognizes specifically for its cognate T7 promoter, which is tightly regulated in the absence of the polymerase (Segall-Shapiro et al., 2014). Hence, it would be advantageous to enable direct thermal control into T7RNAP. To this end, one early attempt had introduced temperature-sensitive intein into T7RNAP gene (Liang *et al*., 2007). The intein was autonomously spliced from the T7RNAP gene at the permissive temperature (18 °C) to enable expressions of both active T7RNAP and associated T7 promoter flanked LacZ product downstream. However, the system was activated at a low temperature (18 °C) which is non-optimal for cell growth and biocatalytic processes, likely experience delays (in hours) due to slow splicing reactions (Shah and Muir, 2014) and non-reversible by nature.

Hence, in this paper, we developed a library of compact, reversible and tunable thermal-repressible split-T7RNAP systems (Thermal-T7RNAPs) which function in the biotechnology-relevant range of 30 – 42 °C. To this end, we leveraged the temperature-sensitive coiled-coil domain of the Tlpa protein as the fusion partner for the N-terminal (NT7) and C-terminal (CT7) based T7RNAP protein fragments, thus enabling direct thermal control of T7RNAP activity. The results indicate that the coiled-coil domain could enable the sequestering of the NT7 and CT7 fusion-fragments at high temperatures (> 40 °C) to inhibit the phage polymerase’s transcription initiation, achieving a high degree of repressibility. The performance of the Thermal-T7RNAPs can be tuned by varying the expression levels and circuit configurations, in which our modelling results revealed that the balance and abundance of the individual protein fragments and its reconstituted form could greatly affect the maximum expression of the GOI and fold change. Furthermore, we also demonstrated the responsive switching kinetics of the Thermal-T7RNAP systems, which was well recapitulated by our model when subjected to dynamic cooling/heating regimes. Next, to achieve temperature tunability as different thermal applications require specific activation temperature, we generated a large mutant library with varying performances and temperature functional ranges via our automated screening and sorting framework. To demonstrate utility, we showed precise temperature control of cell distributions within a microbial co-culture by coupling the developed Thermal-T7RNAP systems with growth inhibition module which regulates glucose uptake. The mutant library provided greater tunability in controlling specific cell distributions. In the process, the developed kinetic model enables the prediction of the growth profiles of individual cell populations within the co-culture mixture. Taken together, we demonstrated the high repressibility, responsive thermal switching kinetics, sharp thermal transition and tunability of our library of Thermal-T7RNAP systems with potential in performing practical applications in biotechnology.

## RESULTS

### Direct thermal control of a heat-repressible split-T7 RNA polymerase (Thermal-T7RNAP)

We hypothesized that direct thermal control over the T7RNAP can be established (Figure 1a) by leveraging the temperature-sensitive C-terminal coiled-coil domain of the Tlpa protein as its fusion partners, which exhibited temperature dependent uncoiling at high temperatures (> 37 °C) (Gal-Mor et al., 2006; Piraner *et al*., 2017). This thermal characteristic would be valuable in bringing apart the N-terminal and C-terminal fragments of split fusion proteins at high temperatures (> 40 °C), and after coiled-coil dimerization, be brought together at lower temperature of 30 °C (Figure 1b). Recent studies have investigated ideal split sites of T7RNAP that allow modification with extra regulatory protein domains and one promising site is the 563/564 position which retained the activity of the polymerase (Baumschlager et al., 2017; Han et al., 2017). Guided by which, we adopted the same split site as the position to fuse with the Tlpa’s coiled-coil. To test the hypothesis, construct in which the N-terminal based (NT7-Tlpa coil) and C-terminal based (CT7-Tlpa coil) fusion proteins were driven by constitutive promoters (gabDP2 and J23101 respectively) was first designed and built to form the Thermal-T7RNAP(v1) system (Supplementary Figure S1a).

**Figure 1.**
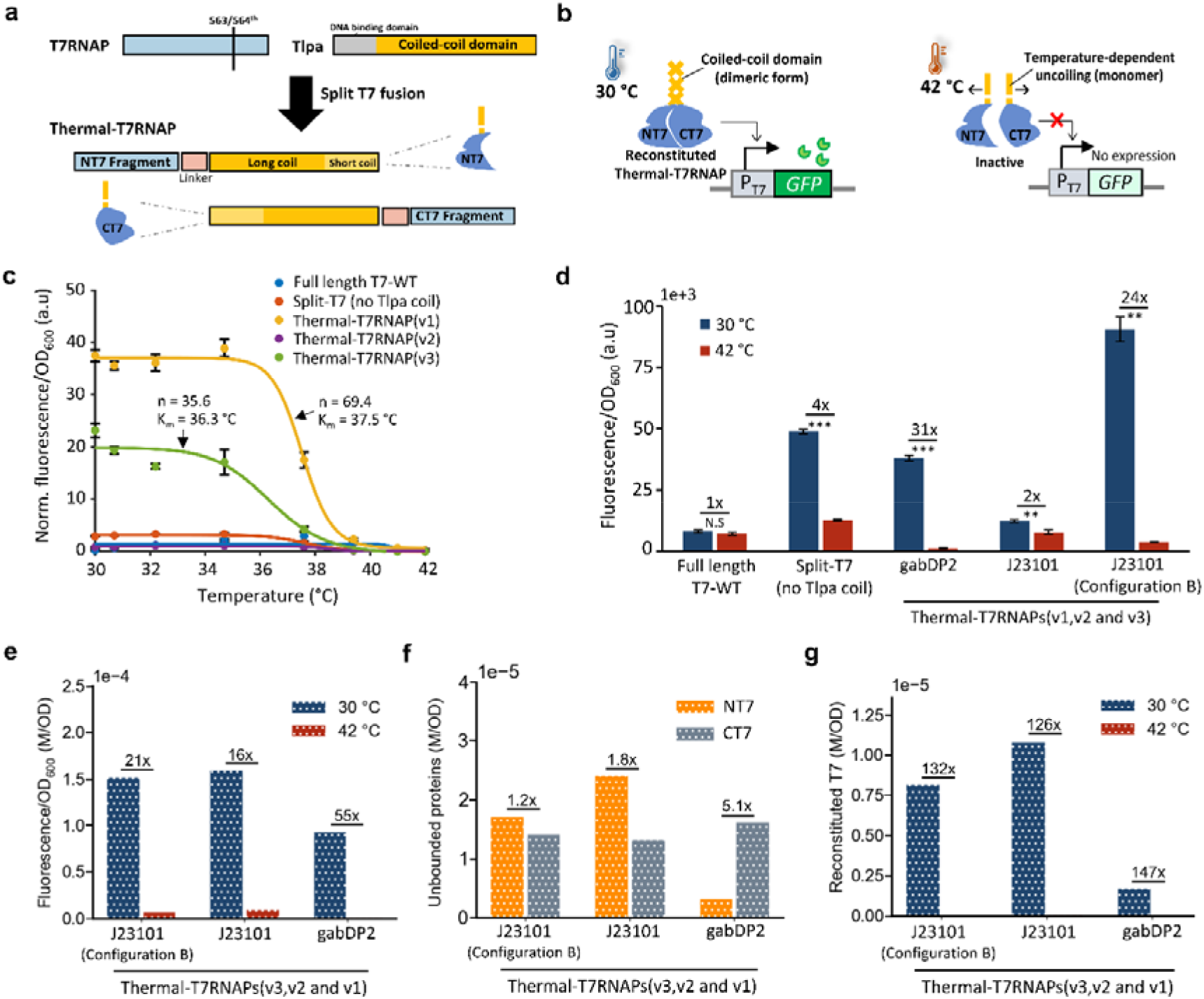
Direct thermal control of Thermal-T7RNAP. **(a) Engineering strategy.** The NT7 and CT7 protein fragments are fused to the temperature-sensitive coiled-coil domain of the Tlpa protein, via a short linker. The coiled-coil domain consists of the longer coiled-coil (94-257^th^) and the shorter coiled-coil (258-371^th^) regions. **(b) Thermal-T7RNAP binding interactions at different temperatures**. At high temperatures, the Tlpa coiled-coil would undergo temperature-dependent uncoiling and allow sequestering of the NT7 and CT7 domains, thus rendering the polymerase inactive for downstream expression to occur. At low temperature, the coiled-coil would retain a dimer complex, allowing the NT7 and CT7 domains to be reconstituted into an active polymerase and capable of driving downstream gene expression. **(c) Temperature dose-response curve**. After an 18-hour incubation period in the thermal regulation assay, the corresponding experimental data (GFP/OD_600_) (filled circles) were normalised to the lowest values at 42 °C and plotted. The dose responses were represented by the Hill equation which were indicated as solid lines. The Hill coefficient, n and half-activation temperature, K_m_, are indicated for some systems (Supplementary Table S1). **(d) Corresponding fluorescence (GFP/OD**_**600**_**) levels after 18 hours**. Constitutive promoters of varying strengths were used in control of the expression levels of NT7-Tlpa coil protein, while the CT7-Tlpa coil protein was being expressed at a constant level using J23101 promoter on another plasmid of similar copy number. An alternative gene configuration B was also adopted to express both proteins on the same plasmid under strong J23101 promoter. The genetic circuits are shown in Supplementary Figure S1. **(e) Predicted fluorescence expression for the Thermal-T7RNAPs after 18 hours. (f) The predicted amount of unbounded NT7 and CT7 fusion proteins at 30 °C. (g) The corresponding predicted amounts of reconstituted T7RNAP proteins at 30 °C and 42 °C**. Data information: The experimental data are represented as mean ± S.D. (n = 3). Statistical significances of ***P < 0.001 and **P<0.01 were calculated based on two-sample unpaired t-test. The corresponding fold-change between the temperatures or the split proteins were shown above each of the bar charts.

We first characterised the Thermal-T7RNAP(v1) system using the thermal regulation assay to study its performance. Within a biological relevant temperature range (30 - 42 °C) that is suitable for cell growth and for many biocatalytic processes, the thermal system exhibited a dynamic range of 31-fold between the permissive (30 °C) and restrictive (42 °C) states and displayed sharp thermal transition centred at 37.5 °C within a narrow functional range of 5 °C (Figure 1c). The temperature response profile was well represented by a Hill equation (Figure 1c and Supplementary Table S1) and revealed the system’s tightness with a maximum repression capacity, K_inh_, of 96 % and a half-activation temperature, K_m_, of 37.5 °C which is defined as the temperature at which the fluorescence intensity is reduced to 50% of its maximum. In comparison, the fold-difference between 30 °C and 42 °C for the Split-T7 system (Figure 1d), which expresses NT7 and CT7 protein fragments solely in the absence of the coiled-coil domains, remains low (4-fold). Importantly, its maximum repression capacity of 75 % was also lower than the Thermal-T7RNAP(v1) system (96%) (Supplementary Table S1), with the basal expression of the Split-T7 system being much higher than that of the Thermal-T7RNAP(v1) at 42 °C (Figure 1d). This suggests that the Tlpa’s coiled-coil domain in Thermal-T7RNAP(v1) is sufficiently strong enough to sequester the CT7 and NT7 protein fragments away from each other and provide high amount of thermal repressibility at high temperatures. In addition, when NT7-Tlpa coil was expressed in the absence of its counterpart CT7-Tlpa coil, it resulted in very minimal expression of the reporter (Supplementary Figure S2). This shows that the high activity of the split polymerase system requires the presence of both protein fragments.

Next, when compared to Thermal-T7RNAP(v1), the full length T7-WT control exhibited 5 times lower maximal expression and no visible fold-difference (Figure 1d). The corresponding growth rate of the full length T7-WT was also 30% lower across the temperature range (Supplementary Figure S3). This may indicate metabolic burden and stress caused by the transcriptionally overactive wild-type polymerase (Temme et al., 2012). We also investigated the use of the mutant T7RNAP(R632S), a variant that was previously adopted in the light inducible split-T7 systems (Baumschlager *et al*., 2017). However, there was minimal expression (data not shown). Thus, we chose the wild-type T7RNAP-WT polymerase as the fusion partners in the Thermal-T7RNAP systems.

In the process, we sought to vary the expression levels and the ratios between the individual NT7-Tlpa coil and CT7-Tlpa coil proteins (Figure 1d) and study how these changes affect the system’s performance. Hence, we constructed Thermal-T7RNAP(v2) by replacing the constitutive promoter that controls the expression level of the NT7-Tlpa coil in Thermal-T7RNAP(v1) from gabDP2 (low) to J23101 (high), while keeping the CT7-Tlpa coil protein expression constant using J23101 on another plasmid of similar copy number (Supplementary Figure S1a). To test a different gene configuration, we created Thermal-T7RNAP(v3) by expressing NT7 and CT7 protein fragments on the same plasmid with the strong promoter J23101 (Supplementary Figure S1b). To gain insights into the different system behaviours, we developed a mechanistic model (Supplementary Table S2) that accounted for the strength of the promoters, the plasmid copy numbers, and importantly, the impact of actual amino acid lengths (NT7-Tlpa coil and CT7-Tlpa coil) on mRNA and protein synthesis rates (Supplementary Figure S5a,b). The model was initially used to capture the behaviours of the Thermal-T7RNAP(v3) at 30 °C and 37 °C (Supplementary Figure S4). To enable the model to predict the performances of different gene circuit configurations at 30 °C (permissive state) and 42 °C (restrictive state), we further incorporated the system’s dose/temperature response (Figure 1c). As a result, the model was able to predict the fluorescent protein expressions, the amount of reconstituted T7RNAP that contributes to fluorescence expression and the abundance of unbounded NT7 and CT7 protein fragments which represent inactive polymerases at the different temperatures (Figure 1e-g respectively).

In general, the model has well predicted the fold-change of Thermal-T7RNAP(v3) fluorescence expression at 30 °C and 42 °C. The relative expression levels of Thermal-T7RNAP(v3) and Thermal-T7RNAP(v1) (Figure 1e) simulated by the model coincided with experimental fluorescence levels (Figure 1d). Using the weak promoter gabDP2 to drive the NT7-Tlpa coil has resulted in the greatest fold-change (31-fold) among the systems which also corresponds with the model prediction showing highest fold-change for the respective promoter (Figure 1d,e). By expressing NT7 and CT7 protein fragments on the same plasmid, the Thermal-T7RNAP(v3) exhibited a comparable fold-difference (24-fold) between 30 °C and 42 °C but achieved two times higher maximum expression, compared to Thermal-T7RNAP(v1) and also captured by the model. The system’s tightness of Thermal-T7RNAP(v3) was also comparable to Thermal-T7RNAP(v1) with a maximum repression capacity of 97% and half-activation temperature of 36.3 °C (Figure 1c and Supplementary Table S1). Although the Thermal-T7RNAP(v2) was predicted to produce fluorescence expression at levels similar to the Thermal-T7RNAP(v3) system (Figure 1e), experimentally (Figure 1d), the system suffered from low expression and fold change, and exhibited considerable leakiness as indicated by the much lower maximum repression capacity of 43%.

The high expression levels of Thermal-T7RNAP(v3) at 30 °C could be due to better balance between the amount of NT7 and CT7 protein fragments as observed in the model (Figure 1f). In comparison, the model revealed a five times lower abundance of NT7-Tlpa coil proteins within the Thermal-T7RNAP(v1) as driven by the weaker gabDP2 promoter (Figure 1f), which limits the total amount of reconstituted T7RNAP to be formed at 30 °C and likely accounted for the roughly 50% lower fluorescence expression observed in experiment and from the model (Figure 1d,e). On the other hand, the slightly higher fold change of Thermal-T7RNAP(v1) compared to Thermal-T7RNAP(v3) could be because the amount of reconstituted T7RNAP generated is within the sensitive region of the T7 promoter (Figure 1g and Supplementary Figure S5c), thus making the system more sensitive to temperature changes. Lastly, the observed lower fluorescence expression than predicted of Thermal-T7RNAP(v2) (Figure 1d,e), where the NT7-Tlpa coil and CT7-Tlpa coil are expressed on different plasmids by the strong J23101 promoters, could be attributed to the excessive accumulation of the NT7-Tlpa coil and reconstituted T7RNAP by more than 30% in comparison to Thermal-T7RNAP(v3) (Figure 1f,g), ultimately affecting its system performance. Taken together, while Thermal-T7RNAP(v1) yielded the highest fold-difference, we also identified Thermal-T7RNAP(v3) which yielded the greatest maximal expression that has potential utility in applications such as bioproduction.

### Dynamic control of Thermal-T7RNAPs and their activation/deactivation kinetics

A key feature of an ideal thermal-switchable system is the ease of tuning the gene expression levels by means of adjustment to the temperature protocols. The dynamic study has two main objectives – firstly, to study whether temperature-controlled (re)activations of existing and *de novo* polymerases can generate reversible gene expression following different cooling and heating regimes and secondly, to leverage on the developed mechanistic model to offer quantitative insights into the inherent kinetics of the Thermal-T7RNAP.

We first investigated the effects of administering a patterned 3-hour (h)-3h-2h-2h OFF-ON (40 - 30 °C) thermal duty cycle, through examining the GFP expressions (Figure 2a) and the associated GFP synthesis rates (Figure 2b) in the Thermal-T7RNAP(v1) and Thermal-T7RNAP(v3) systems. Both systems exhibited similar trends of increase and decrease in GFP expressions at the respective ON and OFF states (Figure 2a). During the ON state (180 – 360 mins), there was a sharp increase in GFP synthesis rates for the initial 60 mins (Figure 2b), which was more apparent for the Thermal-T7RNAP(v3), before attaining the steady synthesis rates in both systems. There was an overall increase in GFP expressions which was mediated by systems’ de-repression when temperature was lowered from 40 °C to 30 °C. In the subsequent OFF state (360 – 500 mins), when the temperature increased to 40 °C, there was an almost immediate repression of GFP synthesis and cessation of GFP accumulation as indicated by the plateau features within the GFP expression profiles (Figure 2a). Noticeably, the transient thermal regulations were highly reversible in both systems and remained sustainable over the full 10h duration. Albeit the lower basal leakiness observed in the Thermal-T7RNAP(v1) system (Figure 2a), the final expression level of the Thermal-T7RNAP(v3) at the end of the regime was much higher than the former.

**Figure 2.**
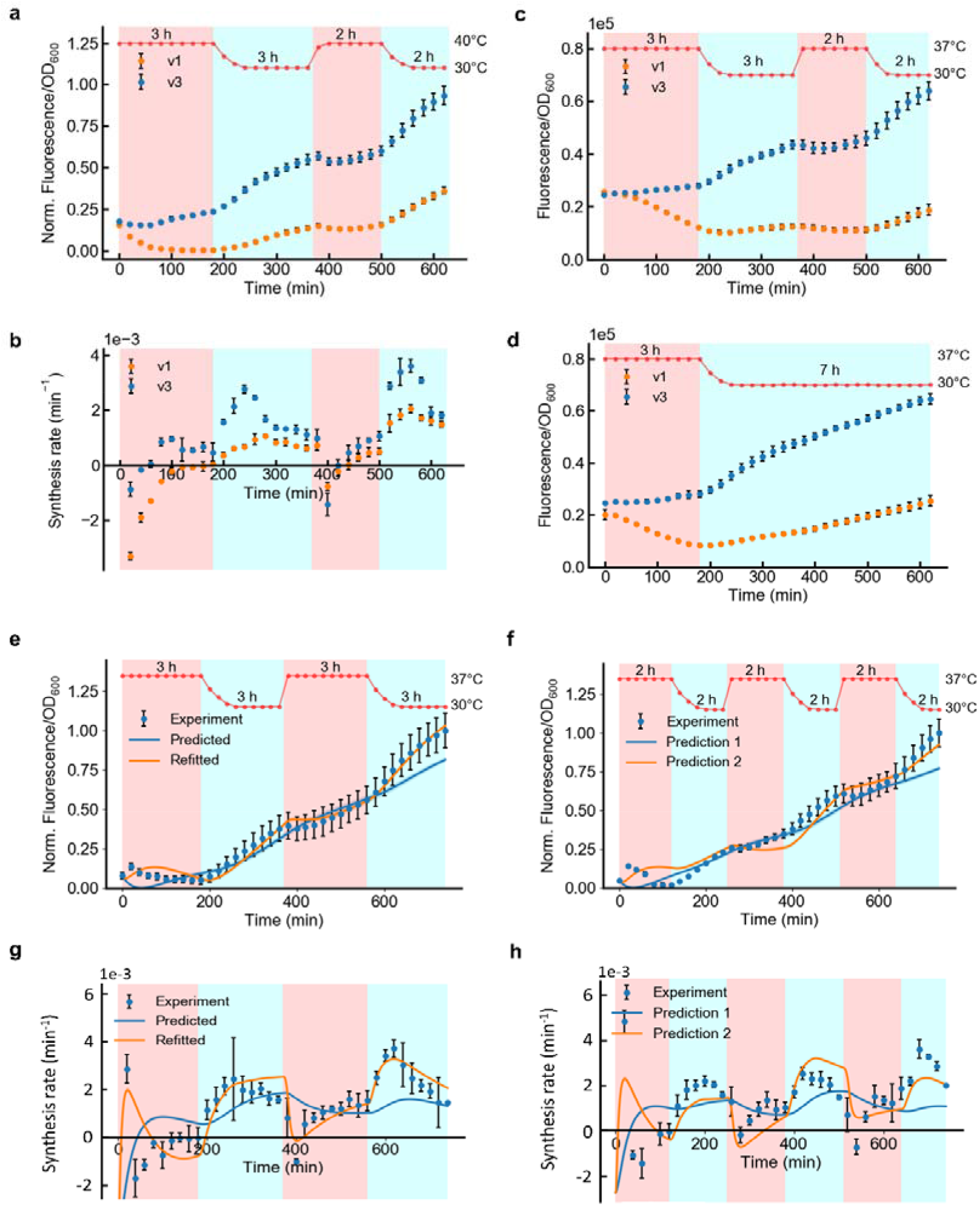
Dynamic control of Thermal-T7RNAPs. **(a) Fluorescence expression profile for Thermal-T7RNAP(v1) and Thermal-T7RNAP(v3) under patterned 3-hour (h)-3h-2h-2h OFF-ON thermal cycle, when alternating between 40 °C and 30 °C.** Measured temperature profiles over the 10-hour period are indicated (red line). Values were normalised to the maximal GFP expression levels. **(b) Corresponding GFP synthesis rates are shown. Absolute fluorescence expression profiles of both systems under (c) patterned 3h-3h-2h-2h thermal cycle and (d) single-step 3h-7h, when alternating between 37 °C and 30 °C**. The absolute values enable ease of comparison of maximum expression levels between systems across the different time interval patterns. **Fluorescence expression profiles for Thermal-T7RNAP(v3) under constant (e) 3h and (f) 2h intervals, when alternating between 37 °C and 30 °C, over 12-hour duration**. The experiment data are shown as filled circles. The solid blue lines represented the predicted profiles derived from a constant temperature regime. The solid orange lines represented the improved predicted profiles after model-refitting and adjustment in model parameters. Their respective growth rates are shown in Supplementary Figure S10. **Corresponding GFP synthesis rates of Thermal-T7RNAP(v3) under constant (g) 3h and (h) 2h intervals are also shown**. Data information: the experimental data are represented as mean ± S.D. (n = 3).

Next, we studied the performance of Thermal-T7RNAP(v1) and Thermal-T7RNAP(v3) systems when the temperatures were altered between 37 °C and 30 °C in the patterned 3h-3h-2h-2h thermal duty cycle (Figure 2c) or within a single-step 3h-7h OFF-ON protocol (Figure 2d). These temperatures are used in conventional practices in bioproduction (Dunstan et al., 2020; Zhang et al., 2018). When compared to previous (40 – 30 °C) thermal cycle study, the systems were not as fully repressed which resulted in slight basal leakiness when the lower OFF state temperature of 37 °C was applied. Nevertheless, despite the difference in time intervals, both systems were still able to achieve highly apparent transient thermal regulations in the 10h duration and even attained comparable final GFP expression values.

We then sought to gain insights on the activation and deactivation kinetics of the systems under the temperature cycling between permissible ON temperature (30 °C) and restrictive OFF temperature (40 °C). While *in vitro* measurements can directly probe into the coiling/uncoiling of alpha-helix structures (Naik et al., 2001; Piraner et al., 2019) in assisting split-polymerase reconstitution/sequestration but precise and reliable measurements of Thermal-T7RNAP initiating transcription are nonetheless undermined by reporter mRNA instability and slow reporter protein turnover (Motta-Mena et al., 2014). As an alternative, we have adopted a mathematical model from earlier work, which has been previously used to provide information of transcriptional kinetics based on activation/deactivation measurements (Motta-Mena *et al*., 2014), to obtain similar transcriptional kinetics information of the Thermal-T7RNAP systems during activation (30 °C) and deactivation phases (40 °C). From the model (Supplementary Figure S6), under the specific temperature regime, the activation (*τ*_*ON*_) and deactivation (*τ*_*OFF*_) time constants represent the respective time taken to attain half-activation of the maximum steady synthesis rate and attain exponential decay to 1/e of its initial rate, as derived from the synthesis rate profiles of the 3h-3h-2h-2h OFF-ON cycle (Figure 2b). Correspondingly, the model was able to recapitulate the dynamic behaviour of the synthesis rates (Supplementary Figure S6c,d). The respective estimated time constants for Thermal-T7RNAP(v1) (*τ*_*ON*_ :24 min;*τ*_*OFF*_ :16 min) and Thermal-T7RNAP(v3) (*τ*_*ON*_:12 min; *τ*_*OFF*_ :14 min) systems fall in the order of tens of minutes. While the deactivation kinetics remained comparable in both systems, it was revealed that the activation kinetics of the Themal-T7RNAP(v3) system was twice as fast as the Themal-T7RNAP(v1) system, suggesting the faster dynamism of the former system to be activated in response to temperature perturbations.

Thus, we further examined the dynamic performance of Thermal-T7RNAP(v3) system under a temperature range commonly used in bioproduction (30 - 37 °C). The intent is to study the induction time response and the performance under different thermal duty cycles (Figure 2e-h). Within the 3-hour time interval OFF-ON cycle which spanned 12 hours (Figure 2e,g), during the ON state (180 – 360 mins), an increase in GFP expression was observed as mediated by system’s de-repression. In the subsequent OFF state (360 – 540 mins), when the temperature increased from 30 °C to 37 °C, there was an immediate repression of GFP expression. In a separate independent 2-hour interval OFF-ON cycle study, similar patterns of GFP expression and protein synthesis rate profiles were also observed and remained thermally responsive despite a reduction in the duration of time intervals (Figure 2f,h).

We then developed a mechanistic model for Thermal-T7RNAP (v3) to gain quantitative insights of its thermal-switching behavior and enable prediction of the thermal performance under different thermal duty cycles. We leveraged the earlier developed mechanistic model (Supplementary Table S2) in predicting the dynamic behaviour of the 3-hour OFF-ON cycle. Generally, when the system was switched to the ON state (Figure 2e), our model revealed an immediate rise in reporter protein synthesis rate prior to attaining its steady rate (Figure 2g). When the system was switched OFF, the model indicated rapid decay in protein synthesis rate and temporary cessation of the total protein level. However, the model (blue line) has underestimated the fast transition kinetics of the experimental dynamic profiles (Figure 2e,g). Interestingly, after increasing the kinetic rates describing the binding-unbinding reaction of the T7RNAP proteins, the performance of the model (orange line) has remarkably improved. This implies that the Thermal-T7RNAP system exhibited fast and sharp binding/unbinding kinetics, with minimum delay in response to alternating temperatures (Figure 2g). This could be due to the fast uncoiling of the temperature-sensitive domains in sequestering the T7 protein fragments at 37 °C, while allowing them to be brought together at 30 °C and in effect, manifested as distinct sharp spikes/dips in individual T7 protein fragments (inactive form) or reconstituted T7RNAP (active form) between the thermal transitions (Supplementary Figure S7). Based on the adjusted model parameters, the T7 transcription rate was also 25% higher, possibly due to better resource allocation when the system was dynamically regulated as opposed to constant induction (Supplementary Table S3). In addition, the binding rate, K_b_ (bounded T7RNAP) and unbinding rate, K_ub_ (unbounded T7RNAP fragments) have increased by 30-fold. Consequently, the 50% lower association constant (K_b_/K_ub_) implies the accumulation of unbounded proteins over its reconstituted form which is likely to account for the improvement in system’s sensitivity to temperature changes. For further validation, the adjusted model (orange line) has also demonstrated robustness in predicting the dynamic behaviour of the independent 2-hour OFF-ON cycle (Figure 2f,h and Supplementary Figure S9).

Using the synthesis rate profiles of 3-hour and 2-hour OFF-ON cycles, the *τ*_*ON*_ and *τ*_*OFF*_ time constants were also derived when temperatures alternated between 37 °C and 30 °C (Supplementary Figure S10). A least-squared error analysis (Motta-Mena *et al*., 2014) was performed to examine the best combination of *τ*_*ON*_ and *τ*_*OFF*_ time constants while cross-referencing with experimental data from both thermal cycles (Supplementary Figure S10c). Under this set of temperature condition and time interval patterns, the analysis show that the estimated time constants of Thermal-T7RNAP(v3) fall within a narrow range (*τ*_*ON*_ :18 - 23 min and *τ*_*OFF*_ :14 - 29 min) with a small margin of error (2 – 3e^-6^).

Taken together, the results suggest that the Thermal-T7RNAP systems exhibited responsive and reversible ON/OFF kinetics following dynamic periods of cooling and heating.

### Automated screening of Thermal-T7RNAP mutants with different functional temperature ranges

Different microbial applications require unique temperature ranges to achieve optimal activities while maintaining the system’s performance. To tune the performance of the Thermal-T7RNAP and create mutants with different thermal characteristics, we mutated the coiled-coil domain of the NT7 and CT7 fusion proteins (Supplementary Figure S11a) which directly influence the GFP expression at the desired repressive temperatures. In this study, we have chosen the Thermal-T7RNAP(v1) as the parent backbone for ease of creating mutants (when the NT7 and CT7 fusion proteins are on separate plasmids) and more importantly this system has the highest fold change out of all the Thermal-T7RNAPs. To facilitate screening of the large library of mutants, we developed an automated screening method for the sorting and identification of ideal temperature-sensitive mutants based on the fluorescence emitted from each colony (Figure 3a). Initially, in the high throughput screening phase, each Thermal-T7RNAP mutant was carefully ‘machine-picked’ with a colony picker and replicative-plated onto three identical 384-format agar plates which were then incubated at the various temperatures (30 °C, 37 °C and 40 °C). Captured information of their overnight fluorescence outputs at respective temperatures was utilised by our in-house data-processing algorithm to sort and identify potential high-performance mutants with desirable fold differences between the permissive (30 °C) and restrictive temperatures (37 °C and 40 °C). Potential candidates were furthered characterised using liquid-cultures over the full range of temperatures (30 – 42 °C). In the process, their individual thermal profiles were automatically fitted with Hill equations and ranked according to their thermal performance parameters such as the half-activation temperature, K_m_, the sharpness of the thermal transition as represented by T_10%_ - T_90%_, the temperature difference (T) between 10% and 90% of maximal fluorescence expression, the associated fold change (Fold) and leakiness (Leak) at various temperatures.

**Figure 3.**
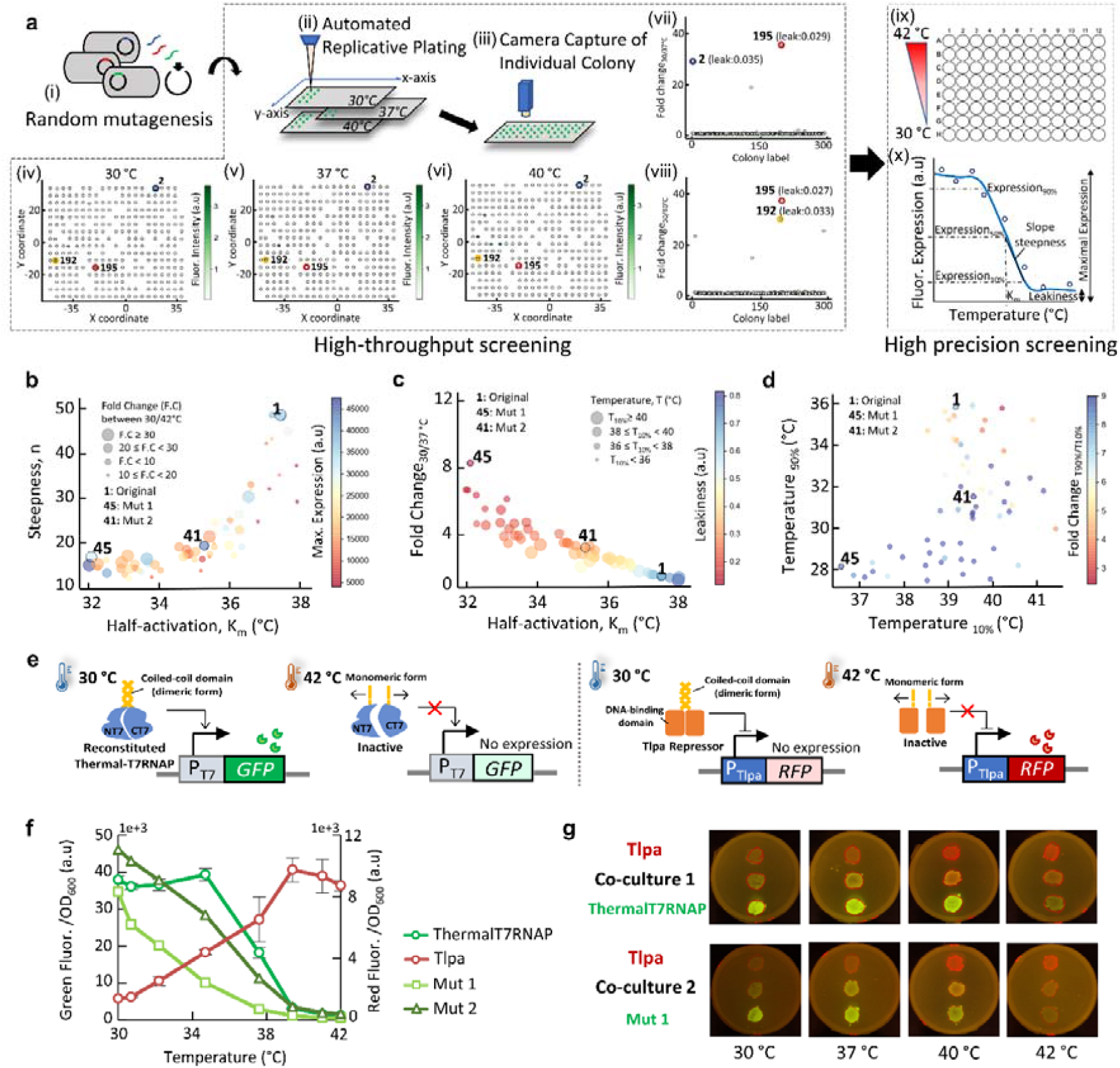
Automated screening of Thermal-T7RNAP mutants. **(a) Mutagenesis framework.** In the first phase of the screening, (i) error-prone PCR was performed on the temperature-sensitive coiled-coil domains of the NT7 and CT7 protein fragments. (ii) The mutant libraries were subsequently transformed and plated on agar plates. The individual overnight colonies were machine ‘picked’ by the colony picker and replicative plated on three identical agar plates for incubation at the desired ‘ON’ and ‘OFF’ state temperatures. (iii) Information of fluorescence and size of each colony was captured by the colony picker (iv-vi) within the agar plates at various temperature. The fold differences in fluorescence between (vii) 30°C and 37°C and (viii) 30°C and 40°C were computed to sort and identify potential mutants that exhibited the desired thermal characteristics. As an example, three potential candidates were highlighted. In the second phase, a high precision screening was conducted. (ix) Liquid characterisation of the shortlisted colonies was conducted in the thermal regulation assay. (x) Each of the dose response was automatically fitted with Hill equation to extract important characteristics. **(b-d) Thermal characteristics of the mutants**. Two discovered mutants (Mut 1 and Mut 2) with distinctively different thermal activation ranges were highlighted. The Temperature _10%_ and Temperature _90%_ represent the temperatures at which the fluorescence levels are at 10% and 90% of their maximal fluorescence expressions respectively. **(e) Mechanisms for the heat-repressible Thermal-T7RNAP system (left) and heat-inducible Tlpa system (right)**. At low temperature (30 °C), the coiled-coil domain of the Tlpa repressor protein maintained as a dimeric complex which allows the binding of the DNA binding domain to its promoter pTlpa to inhibit downstream repression. At high temperatures (> 37 °C), the coiled-coil domains undergo temperature-dependent uncoiling which render the Tlpa repressor inactive, thus enabling of constitutive expression of the promoter. **(f) Thermal dose responses** after 18 hours incubation. **(g) ‘Traffic light’ pattern**. For the co-cultures, the respective cells were grown at equal initial proportions and plated on agar overnight to observe thermal logic. Data information: The experimental data are represented as mean ± S.D. (n = 3).

Using the high throughput system, we screened over 1700 colonies and many exhibited constitutive expressions regardless of temperature change while others exhibited loss of expression (Supplementary Figure S11b). We discovered that mutating the NT7 fusion protein only or simultaneously mutating both the CT7 and NT7 fusion pair yielded very few transformants (∼ 200) and did not exhibit desired thermal characteristics. Nonetheless, we successfully isolated 130 CT7 mutants which exhibited thermal repressibility in the desired temperature range and subjected them to the liquid-culture characterisations of their temperature profiles in the second screening phase (Figure 3a). We identified two mutants (Mut 1 and Mut 2) that retained the ideal characteristic of the original Thermal-T7RNAP(v1) system but with distinctive shift in their temperature functional ranges (K_m, original_= 37.5 °C, K_m, Mut1_ = 32.2 °C and K_m, Mut2_ = 35.4 °C) (Figure 3b). The Mut 2 system also exhibited 20% higher maximal expression when compared to the original system. While both mutants had more gentle thermal transitions ([T_10%_ - T_90%_]_original_ = 3.3 °C, [T_10%_ - T_90%_]_Mut1_ = 8.5 °C and [T_10%_ - T_90%_]_Mut2_ = 8.0 °C) (Figure 3d), lower leakiness were exhibited by the mutants at 37 °C ([Leak_30°C - 37°C_]_original_ = 0.68, [Leak_30°C - 37°C_]_Mut1_ = 0.12 and [Leak_30°C - 37°C_]_Mut2_ = 0.31) (Figure 3c). Correspondingly, greater fold differences between 30 °C and 37 °C in Mut 1 and Mut 2 were observed ([Fold_30°C - 37°C_]_original_ = 1.50, [Fold_30°C - 37°C_]_Mut1_ = 8.53 and [Fold_30°C - 37°C_]_Mut2_ = 3.27) (Figure 3c).

The ability of simultaneously thermally activating and inhibiting expressions in different cell populations can be a powerful tool in biotechnology. As a proof of concept, we coupled our heat-repressible Thermal-T7RNAP system to control GFP expression in one cell and the heat-inducible Tlpa system to control RFP expression in another cell (Figure 3e). The thermal logic is illustrated by both thermal systems which functioned in counter-unison to produce higher GFP expressions at low temperatures (< 34 °C) and higher RFP expressions (due to de-repression of Tlpa repressor from promoter pTlpa) at high temperatures (> 38 °C) (Figure 3f). The two Thermal-T7RNAP mutant systems (with smaller K_m_) in the thermal profiles have left-shifted their temperature intersections with the Tlpa system. Similarly, in the ‘Traffic light’ agar patterns (Figure 3g), shifting of the intersection was portrayed by the diminishing intensity of the ‘orange zone’ in the Mut 1/Tlpa co-culture when compared to the original co-culture at higher temperatures (> 40 °C).

### Directing microbial community distribution with thermal control

The use of thermal biosensors in controlling cell distribution within microbial community is still not well undertaken. While existing study leveraged optimal temperature ranges to enable native microbial species to co-exist (Krieger et al., 2021), there was a lack of capability in active control of the individual population. To address the need, we developed temperature-based genetic circuits which forms thermal logic to enable thermal modulation of the growth of two engineered *E. coli* populations within a co-culture (Figure 4a), in which one population is producing GFP while another is producing RFP. We built a thermal-repressible ThermalT7RNAP-SgrS system for the GFP-reporting cell (GFP_ThermalT7RNAP-SgrS_) and a thermal-inducible Tlpa-SgrS system for the RFP-reporting cell (RFP_Tlpa-SgrS_) (Figure 4b and Supplementary Figure S12). Both thermal circuits were designed to slow down cellular growth by limiting glucose uptake at their ‘ON’ thermal states, by expressing the SgrS sRNA (sugar transport related silencing RNA) which functions to degrade ptsG mRNA that encodes major glucose transporter, IICB^Glc^ (Negrete et al., 2010; Wadler and Vanderpool, 2007).

**Figure 4.**
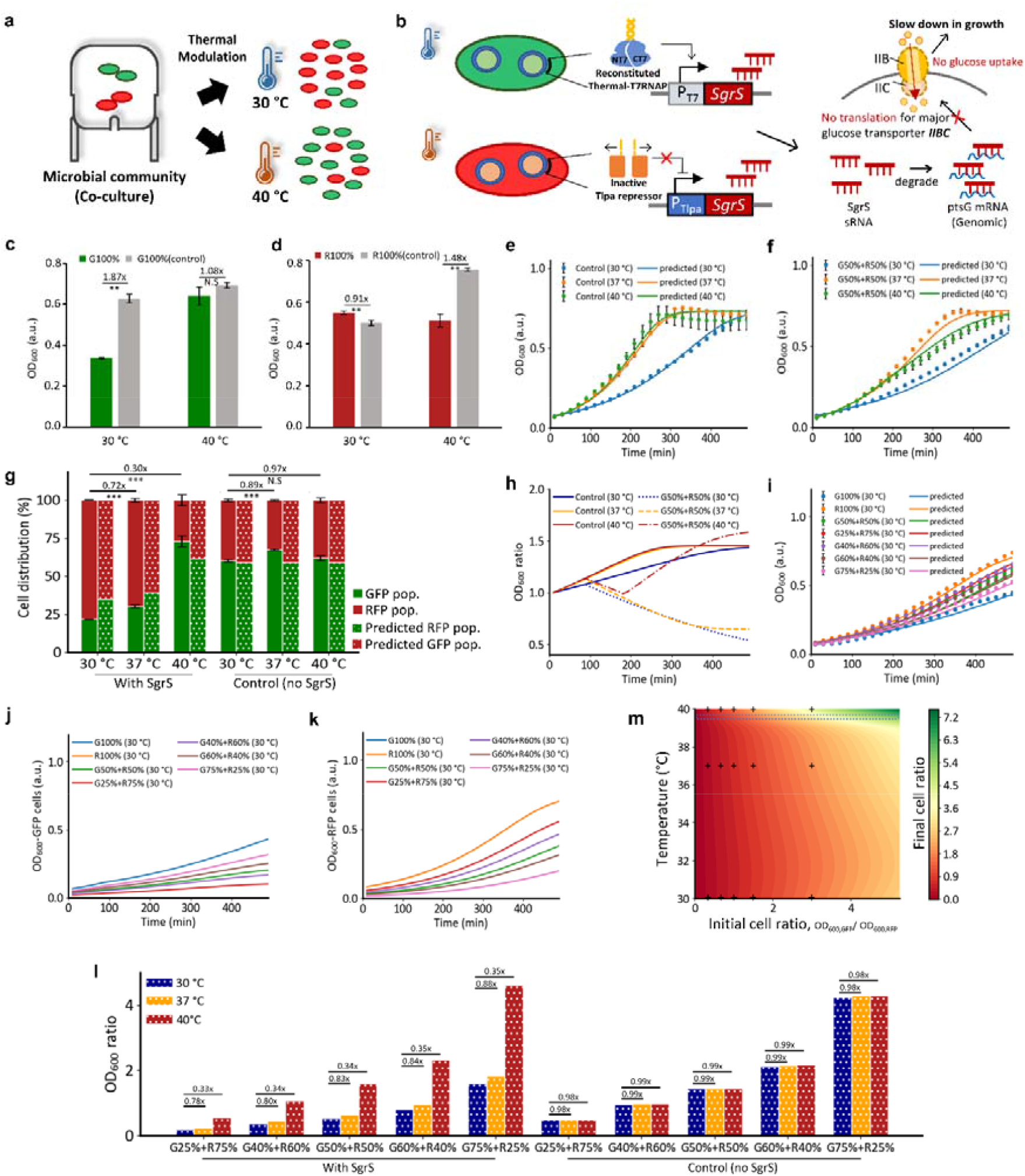
Thermal control of co-culture distribution. **(a) Thermal modulation.** At different temperatures, the cell proportions are regulated. **(b) Growth inhibition mechanisms**. The GFP reporting cells contained the heat-repressible Thermal-T7RNAP system and the RFP reporting cells contained the heat-inducible Tlpa system. Both systems regulate their cell growth by limiting glucose uptake at 30 °C and 40 °C respectively. **(c**,**d) Final growth values for the monocultures (100%) at 30 °C and 40 °C after 8 hours**. The GFP (G) and RFP (R) cells which contained the growth inhibition modules and their respective controls (no SgrS expression) were shown. **(e**,**f) Combined growth curves for the co-cultures**. The cells were initially seeded at equal proportion (G50%+R50%). The control co-culture does not express SgrS. Predicted growth curve of the co-culture (solid lines) and the experimental data (filled circles) were shown. **(g) Final cell distributions in the co-culture**. After 8 hours, the final compositions between the GFP population and RFP population were derived from their measured fluorescence levels (solid bars). The amount of GFP cells was computed as a percentage of the fluorescence intensities of the monoculture at 40 °C (repressed). The amount of RFP cells was obtained after subtraction from the GFP cells. The shaded bars represented the predicted cell distributions as derived from the ratio between individual ODs. **(h) Change in OD ratios over time**. The cells were initially seeded at equal proportion. **(i) Combined growth curves under various initial cell proportions**. Predicted growth curve of the co-culture (solid lines) and the experimental data (filled circles) were shown. **(j**,**k) Predicted growth profiles of individual GFP and RFP. (l) Final OD ratios of the co-cultures. (m) Contour representation of initial and final cell proportions within the co-culture, under different temperatures**. The cell proportions are obtained by dividing the predicted OD values of GFP cells with the RFP cells. The cross marks represented conditions performed in experiment. The highlighted blue region represented the conditions that maintained cell proportions in the 8-hours period. Data information: The experimental data are represented as mean ± S.D. (n = 3). Statistical significances of ***P < 0.001 were calculated based on two-sample unpaired t-test. The fold change between the temperatures were indicated above each of the bar charts.

We first conducted independent characterisation of the ThermalT7RNAP-SgrS and the Tlpa-SgrS systems to study the growth inhibition profiles at different temperatures (Figure 4c,d and Supplementary Figure S13). At 30 °C, there was an 87% decrease in the growth of the GFP_ThermalT7RNAP-SgrS_ cells based on the final OD values when compared to its corresponding control that had no SgrS (Figure 4c). But at 40 °C, there was no observable growth inhibition, showing the tightness of the system in preventing SgrS expression. Conversely, at 40 °C, the RFP_Tlpa-SgrS_ cells displayed a 48% reduction in growth compared to its control but exhibited normal growth at 30 °C, implying strong repression of SgrS expression at 30 °C (Figure 4d).

In parallel, we developed a growth model to investigate the effects of temperature variations on cell growth and changes in glucose uptake due to the expression of SgrS in the monocultures (Supplementary Figure S16 and Supplementary Table S5-S6). In the model, the maximum growth capacity of the monoculture was presumed to be constrained by the amount of glucose carbon source. The RFP control cells were slower in growth in comparison to GFP control cells with maximum specific growth rates of 0.014 min^-1^ and 0.016 min^-1^ respectively (Supplementary Table S6).

The growth rates of both control cells also increased with increasing temperatures and defined by half-activation temperature of 29.24 °C and steepness of 10.92 (Supplementary Table S6). The glucose sensitivity was estimated by the model to be 0.759 g/L with a yield coefficient of 0.122 (Supplementary Table S6). By introducing SgrS expression at either 30 °C or 40 °C for the GFP_ThermalT7RNAP-SgrS_ and RFP_Tlpa-SgrS_ cells respectively, the effects of their growth inhibition were clearly visible in both cells by an approximately 7 to 8-fold reductions in glucose sensitivity compared to their controls which led to reduced glucose uptake (Supplementary Table S6). We also observed an initial delay (first 200 mins) in glucose uptake inhibition for the RFP_Tlpa-SgrS_ cells but was absent from the GFP_ThermalT7RNAP-SgrS_ cells, which again demonstrated the fast thermal response of the Thermal-T7RNAP in driving the SgrS expression (Supplementary Figure S16).

We then proceeded with the characterisation of the co-culture in which both GFP_ThermalT7RNAP-SgrS_ and RFP_Tlpa-SgrS_ cells were grown to similar initial ODs and mixed at equal proportion (G50%+R50%) (Supplementary Figure S14). We sought to demonstrate that the proportion of the cells can be actively controlled using the thermal genetic circuits (i.e., the final cell distributions can be varied). We used the growth model developed from the monoculture (Supplementary Table S5) to predict the individual growth profiles of the two cell populations within the co-culture (Supplementary Figure S18). The growth of the two cell populations was presumed to be constrained by the shared carbon source. Their combined growth was thus computed as the sum of the individual cell populations. As evidenced, the predicted combined growth profiles of the co-culture (G50%+R50%) at different temperatures agreed well with the experimental results (Figure 4e,f). This implies that the growth model could serve as a means to predict the actual growth profiles of the individual cell population within a co-culture.

While fluorescence-reporting is commonly used to estimate the individual cell growths within a co-culture (Chait et al., 2017; Nikolic et al., 2013; Stephens *et al*., 2019), we observed that the correlations between the reporting fluorescence and OD values were different even for cells with similar genetic makeup (Supplementary Note S1 and Supplementary Figure S17). This indicates that the use of fluorescence reporting level to estimate the individual cell growths within a co-culture may not be a good representative metric. Hence, for better representation, we derived the final cell distributions of the co-cultures under different temperatures from both the experimentally measured fluorescence and the model-predicted individual growth profiles (Figure 4g and Supplementary Figure S18). At 30 °C, the cell distributions for both experiment and prediction have exhibited prominent decline in the GFP_ThermalT7RNAP-SgrS_ population from the initial G50%+R50% to attain final ratios of [G22%+R78%]_Expt._ and [G35%+R65%]_Predict._ respectively (Figure 4g). At 37 °C, the final ratios of [G30%+R70%]_Expt._ and [G39%+R61%]_Predict._ attained were comparable to 30 °C, indicating that the growth of the GFP_ThermalT7RNAP-SgrS_ population was still inhibited. This suggests that there was still significant amount of SgrS expression within the GFP_ThermalT7RNAP-SgrS_ population. It was at 40 °C when GFP_ThermalT7RNAP-SgrS_ cells had outgrown the RFP_Tlpa-SgrS_ cells; to attain final ratios of [G73%+R27%]_Expt._ and [G61%+39%]_Predict._. This reiterated that the ThermalT7RNAP-SgrS system remained tightly repressed at high temperatures which allowed the GFP_ThermalT7RNAP-SgrS_ cells to grow normally while the Tlpa-SgrS system produced the desired slowdown in the growth of RFP_Tlpa-SgrS_ cells. For the control co-culture, the final distributions hovered around [G60%+R40%]_Expt. & Predict.,_ across all the temperatures tested which implied a roughly 10% increase in GFP control cells even though both GFP and RFP control cells were initially cultured at equal proportion.

To reveal changes in cell proportions over time, we computed the OD ratios through dividing the predicted ODs of the GFP cells with the RFP cells (Figure 4h and Supplementary Figure S18). In the control co-culture (Figure 4h), we observed an increase in the OD ratios from 1.0 (equal initial cell proportions) to 1.4 eventually; showing the faster growth rate of GFP control cells. In the co-cultures with the thermal gene circuits (Figure 4h), at 30 °C and 37 °C, there were reductions in the OD ratios over time to finally attain ratios of 0.5 and 0.6 respectively. At 40 °C, there was a slight decrease in the first 200 mins followed by the steady increase in the OD ratio before reaching 1.6. The initial dip in OD ratio was largely due to the mentioned delay in SgrS expression of RFP_Tlpa-SgrS_ cells while its 14% increase in final ratios compared to control co-culture was attributed to the decline in growth of RFP_Tlpa-SgrS_ cells as a result of SgrS expression at high temperatures.

Next, we investigated the various initial cell proportions in affecting the final cell distributions within the co-cultures (Figure 4i-k and Supplementary Figure S15). The fitted growth model of the monocultures successfully predicted their combined growth profiles under different temperatures and (Figure 4i and Supplementary Figure S19-S20). At different initial proportions for control co-cultures, there is a constant 40% increase from their initial ratios (Supplementary Figure S21) and their final cell ratios remained similar at different temperatures (Figure 4l). In the co-cultures with the thermal genetic circuits, by lowering or raising the temperature, we successfully modulated cell ratios as much as -50% and 65% respectively (Supplementary Figure S21). From model, we also identified the fine-tuning conditions required to obtain specific cell proportions (Figure 4m). We discovered that the final cell proportions can effectively be maintained (within a 10% deviation) with respect to their initial proportions at 39.5 °C.

To expand the tunability of thermal modulation in co-culture, we leveraged the discovered Thermal-T7RNAP CT7 mutants (Mut 1 and Mut 2) to regulate the SgrS expression in the GFP cells (Figure 5a-c). In the earlier results, both mutants have distinctive left shifting of activation temperature (Figure 3f) that can potentially provide better repression capability at 37 °C which is a common temperature for cell cultivation.

**Figure 5.**
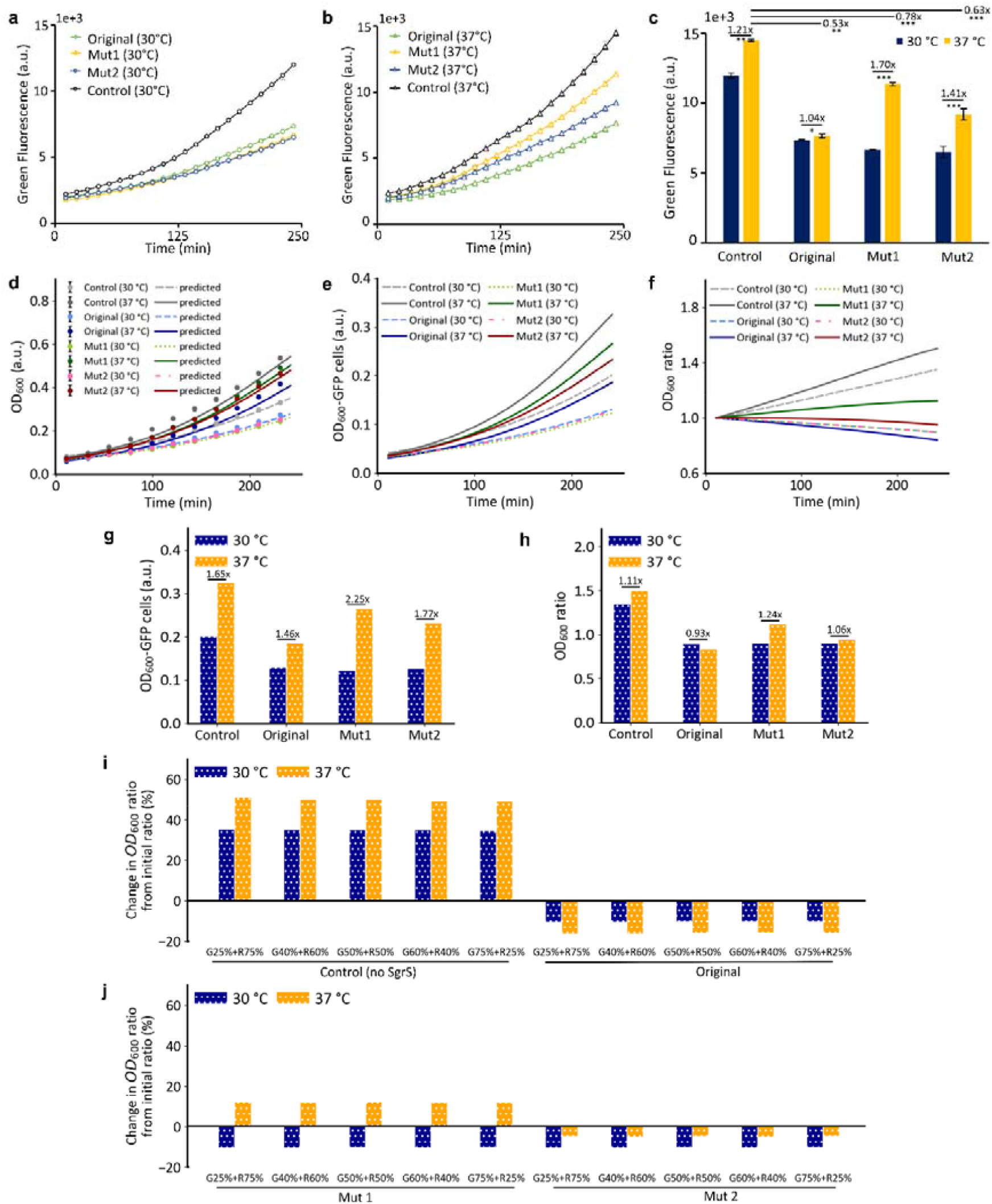
Thermal modulation of co-culture with Thermal-T7RNAP mutants. **(a,b) Continuous fluorescence intensities profiles of the co-culture library at 30 °C and 37 °C.** The GFP_ThermalT7RNAP-SgrS(unmutated),_ GFP_ThermalT7RNAP-SgrS(Mut1)_ and GFP_ThermalT7RNAP-SgrS(Mut2)_ cells were each grown together with the RFP_Tlpa-SgrS_ cells at equal initial proportion to form the original, Mut 1 and Mut 2 co-culture systems respectively. **(c) Corresponding fluorescence values after 250 mins. (d) Combined growth curves of the co-cultures**. The experimental (filled circles) and predicted values (solid lines) were shown. **(e) Predicted OD growth curve for the GFP population. (f) OD ratio profiles over time**. The ratios were computed after diving the predicted OD values of the GFP population by the RFP population. **(g) Predicted OD values for GFP cells** and **(h) final OD ratios** after 250 mins. **(i**,**j) Change in OD ratios between initial and final conditions of the co-cultures under different initial cell proportions** after 250 mins. Data information: The experimental data are represented as mean ± S.D. (n = 3). Statistical significances of ***P < 0.001, **P<0.01 and *P<0.05 were calculated based on two-sample unpaired t-test. The corresponding fold-change between the temperatures were shown above each of the bar charts.

In the original G50%+R50% co-culture, the GFP_ThermalT7RNAP-SgrS_ cells were cultured together with the RFP_Tlpa-SgrS_ cells at equal proportion. Results show that at 30 °C, the GFP_ThermalT7RNAP-SgrS_ cells exhibited the desired growth inhibition as portrayed by the 63% decrease in fluorescence intensity from the GFP control cells (Figure 5a,c). However, at 37 °C, there was still a significant 47% reduction in the fluorescence intensity from the control (Figure 5c) which indicated that growth inhibition was still present at this higher temperature. For Mut 1 and Mut 2 co-cultures which contained the GFP_ThermalT7RNAP-SgrS(Mut1)_ and GFP_ThermalT7RNAP-SgrS(Mut2)_ respectively, at 37°C, the fluorescence reductions from the GFP control cell were lesser (22% and 37% respectively) (Figure 5c), which suggests that the mutant systems exhibited greater relief from SgrS expression at 37 °C while still achieving desirable growth inhibition at 30 °C.

Similarly, we predicted the growth profiles of the mutant co-cultures at different initial cell proportions (Figure 5d,e and Supplementary Figure S22-S24). The relief in SgrS inhibition was also visible from the predicted final OD values of the GFP cells (Figure 5g) where GFP_ThermalT7RNAP-SgrS(Mut1)_ cells achieved 125% increase in cell growth at 37 °C from 30 °C, followed by 77% and 46% increase for GFP_ThermalT7RNAP-SgrS(Mut2)_ and the GFP_ThermalT7RNAP-SgrS_ respectively. We also observed that while the original system exhibited decrease in the OD ratio at 37 °C, the ratios for Mut 1 and Mut 2 systems were maintained over time (Figure 5f,h) and the same is true for the other initial cell proportions (Figure 5i,j).

## DISCUSSION

In the thermogenetic toolkit, existing heat-repressible systems are limited in parts (Hoynes-O’Connor *et al*., 2015; Liang *et al*., 2007; Qing *et al*., 2004; Yang Zheng *et al*., 2019) and restricted to thermal sensing mechanisms which relied mainly on RNA or transcription factors; and faced potential issues such as functioning at low temperatures (< 30 °C) which are non-optimal for cell growth and bio-catalytic processes, have wide temperature transitions, suffer delays or complex in design. To address the limitations and provide an alternative mode of thermal regulation, in this paper, we created a library of novel heat-repressible Thermal-T7 RNA polymerase (Thermal-T7RNAP) systems by fusing Tlpa coiled-coil domain with split-T7RNAP to introduce direct temperature control into the polymerase. The results show that the polymerase activity was well repressed at high temperatures (> 40 °C) and highly suggests the NT7 and CT7 protein fragments can be sequestered by the temperature-sensitive coiled-coil domain at high temperatures (Figure 1). The system has several advantages over existing heat-repressible systems. Firstly, the system is compact and highly simplified in design which encourages ease of incorporation into larger and more complex genetic networks to perform integrative functions. Secondly, the system is responsive and highly reversible, which is useful for transient thermal regulation of gene expressions. Thirdly, the system’s temperature transition is narrow (5 – 8 °C) and functions within a tunable temperature range (30 – 42 °C) which is optimal for cell growth and bio-catalytic processes. Lastly, as the Thermal-T7RNAP system utilised the T7 phage polymerase within its design core, it can decouple transcription from the host’s machinery and presume to be highly suited in performing independent functions within a large group of organisms. As an exemplification of novel thermal application, we demonstrated that *E. coli* cells population distributions in a co-culture can be actively altered using temperature control (Figure 4 and 5).

We have developed model to gain insights into the behaviour of the Thermal-T7RNAP systems under different expression levels of the NT7-Tlpa coil and CT7-Tlpa coil proteins (Figure 1). Differences in performance were observed when we experimented with constitutive promoters of varying strengths and circuit configurations. The model predicted well the fold change of two of the Thermal-T7RNAP systems and their relative maximum expressions. Interestingly, the model revealed that the highest fold change of Thermal-T7RNAP(v1) is likely due to the lower amount of reconstituted T7RNAP expressed (Figure 1f-g) which possibly falls within the sensitive range of the P_T7_ promoter curve (Supplementary Figure S5c). On the other hand, expressing greater amount of both protein fragments but at similar proportions displayed the highest maximal expression of the reporter as seen in Thermal-T7RNAP(v3) (Figure 1d-g).

One unique feature of the Thermal-T7RNAP system is the ability to undergo sharp two-way thermal switching, which is likely attributed to the highly sensitive temperature-dependent coiling and uncoiling of the coiled-coil domain (Figure 2). The model unveiled a beneficial 50% reduction in association constant when the system was regulated dynamically (Supplementary Table S3). This implied a larger pool of unbounded NT7 and CT7 protein fragments would be readily available for reconstitution in the upcoming activation phase and led to the estimated half-activation response within tens of minutes. This suggests that the thermally responsive characteristic of Thermal-T7RNAP system is potentially useful in temperature-based dynamic regulation of bio-processes for example in bioproduction relevant range (30 – 37 °C) such as in the control of metabolic fluxes (Harder et al., 2018; Wang et al., 2021), that can complement with existing light-based systems (Zhao et al., 2019; Zhao et al., 2018). During the activation phase, when the temperature alternated between 30 °C and 40 °C (Supplementary Figure S6c,d), the initial spike in GFP synthesis rate could be attributed to the faster initial increase in cell growth before growth stabilisation (data not shown), although the artefact was not as apparent when alternating between 30 °C and 37 °C. To the best of our knowledge, this is a first characterization of thermal-switchable systems where different cooling and heating regimes were applied to study the dynamic performance of thermal-switchable systems, in which the Thermal-T7RNAPs were observed to be highly reversible while maintaining robust thermal performance. This approach of dynamic characterisation enables us to study the reversibility of the thermal systems. While the activation and deactivation kinetics are highly dependent on the temperature regime, (i.e. thermal transfer gradient), the model-derived estimation of the time constants can nevertheless enable direct comparison to be made between the systems under the same temperature regimes. By fitting thermally-regulated dynamic expression profiles with mechanistic and/or simple kinetic models, it can be highly valuable for bioproduction whereby one can derive the appropriate cooling/heating gradients and the desired duration and strength of thermal induction to achieve eventual performance. Although reasonable fold changes were observed in current systems, it can be further improved to suit specific applications potentially via the addition of positive feedbacks (Nistala et al., 2010) and/or introducing temperature-sensitive proteolytic modules (Yang Zheng *et al*., 2019).

Different applications require specific temperature activation points. Here, we created a library of Thermal-T7RNAPs functioning between 30 – 42 °C. The wild-type Thermal-T7RNAPs which offer better repressibility at high temperatures (K_m_: ∼ 37.5 °C) can potentially function as *in vivo* fever sensor or temperature-actuated kill switch which have much higher temperature requirements (> 40 °C) to avoid non-specific activations (Piraner *et al*., 2017). Whereas, the two identified high-performance mutants (K_m_ < 37.5 °C) which have better repressibility at 37 °C could serve as ideal growth switches (Fang et al., 2020; Harder *et al*., 2018; Schramm et al., 2020) within biocatalytic processes to segregate microbial growth (37 °C) from subsequent bioconversion phase (30 °C). The automated screening and sorting framework for temperature-sensitive mutants has also shortened the overall duration and the need for manual intervention (Figure 3a), as compared to manual screening method (Piraner *et al*., 2017). We discovered solely mutating the temperature-sensitive coiled-coil domain on the CT7 fusion protein while retaining the wild-type NT7 counterpart can create distinctly different thermal profiles of the heat-repressible system (Figure 3b-d). The possible presence of consensus sequences within the coiled-coil domains of wild-type and mutant systems may highlight the likely presence of essential amino acid residues that resided at these specific regions which are vital for proper interfacial ionic interactions that governed the C_2_ symmetry of the parallel coiled structure (Piraner *et al*., 2019) and ensuring their robust thermal performances. Furthermore, the improvement in maximum expression or reduced leakiness could also be due to the formation of heterodimeric coiled-coils as opposed to homodimers formation of the wild-type system, that beneficially omits background noise associated with the NT7-NT7 or CT7-CT7 fusion pairs (Piraner *et al*., 2019).

To extend the use of thermal control in microbial applications, it is useful to enable independent control of multiple functions within a tunable temperature range (Figure 3e-g). To the best of our knowledge, this is the first study to demonstrate the utility of both thermal-repressible and thermal-inducible systems to enable thermal logic in the active control of population dynamics within a microbial community (Figures 4 and 5) that experienced growth disparities among the populations when un-regulated (Figure 4g,h). Using the thermal modulation logic, we effectively altered the cell distribution between the two cell populations of the co-culture by reducing their individual growth rates under different temperature settings (Figure 4g). At the low temperature of 30 °C, with an initial cell proportion of 1:1, we modulated the cell distribution to reach a final ratio of 3:7. At 40 °C, the final ratio can be reversed to 7:3. This is possible as the thermal–repressible system functions at a temperature range suitable for the microbial cell growth.

The use of fluorescence-reporting is a common strategy to estimate the individual cell growths within a co-culture (Chait *et al*., 2017; Nikolic *et al*., 2013; Stephens *et al*., 2019). However, from the monoculture data, we revealed that the correlations between the reporting fluorescence and OD values can be distinctively different at different temperatures even when they carried similar genetic make-ups (Supplementary Note S1). Hence, in this study, we developed a growth model built upon the monoculture data (Supplementary Figure S16), which allows us to better estimate the distribution of individual cell populations within a co-culture mixture (Supplementary Figure S18). The model also provides insights into the initial conditions required to achieve a specific final cell proportion, which enables one to precisely manipulate the ultimate cell distribution (Figure 4m). The extensive Thermal-T7RNAP mutant library (Figure 3) also enabled customisability in varying the levels of growth inhibition within the co-cultures and consequently provided greater flexibility in thermal modulation, for both maintenance and alteration of the cell proportions (Figure 5).

Taken together, the Thermal-T7RNAPs has expanded the synthetic biology thermal control toolkit for biomedical (Abedi et al., 2020; Gamboa et al., 2020; Miller et al., 2018; Piraner *et al*., 2017) and industrial applications (Harder *et al*., 2018; Rodrigues et al., 2017; Salila Vijayalal Mohan et al., 2020; Wang *et al*., 2021; Xu et al., 2020) by providing an alternative mode of thermal regulation. The Thermal-T7RNAPs can be an useful addition for combinatorial multi-level temperature-based gene regulations (e.g. combining transcription, post-transcriptional and translational control) (Greco et al., 2020; Naseri and Koffas, 2020). Potentially, it can be an envisioning tool for controlling the complex inter-population dynamics of microbial consortia in emerging biomedical and bioproduction applications.

## DATA AVAILABILITY

The source data underlying Figures 1 to 5 are provided as a source data excel file upon request. All other relevant data, including plasmid sequences and plasmids that support the findings of this study are additionally available from the corresponding author upon request.

## SUPPLEMENTARY DATA

Supplementary information is available.

## CODE AVAILABILITY

Python 3 (Version 3.7.1) was used to implement all the models. The developed models underlying Figures 1, 2, 4 and 5 are available as source codes in the respective folders upon request.

## ACKNOWLEDGEMENTS

The authors thank Dr. Chong Da Tan from Singer Instruments for providing valuable advice and guidance on the use of the colony picker. The authors also thank Miss Sheena Chan for her advice on the growth inhibition modules.

## AUTHOR CONTRIBUTIONS

W.K.D.C., J.W.Y. and C.L.P. conceived the ideas. W.K.D.C. designed and performed the experiments. All authors analysed the results. J.W.Y. developed and simulated the models. W.K.D.C. and V.L.D. performed the mutagenesis and J.W.Y. processed the data. All authors wrote and edited the manuscript. C.L.P. supervised the project.

## MATERIALS AND METHODS

### Plasmid design and construction

All the plasmids were designed *in silico* using Benchling (Benchling, Inc. San Francisco, CA, USA). Individual gene fragments and primers were synthesized from Integrated DNA Technologies (Integrated Device Technology, Inc. San Rose, CA, USA). The polymerase chain reaction (PCR) products were amplified using Q5 High-Fidelity DNA polymerase (New England Biolabs, MA, USA) with strict accordance to the manufacturer’s protocols. PCR products were analysed by gel electrophoresis using 1% agarose gel and purified using QIAquick gel extraction kit (Qiagen, Hilden, Germany). The DNA concentrations of the gel-purified samples were quantified with Nanodrop One^C^ (Thermo Fisher Scientific, MA, USA). Gibson assembly was subsequently performed using the NEBuilder HiFi DNA assembly (New England Biolabs, MA, USA) with strict accordance to the manufacturer’s protocols. Subsequent assembly products were chemically transformed into *E. coli* K-12 strain NEB *DH-10 Beta* (New England Biolabs, MA, USA) unless stated otherwise. Colonies that grown on the LB-antibiotic plate were picked and inoculated into fresh LB-antibiotic medium at 37 °C to prepare overnight culture for plasmids extraction using QIAprep Spin Miniprep kit (Qiagen, Hilden, Germany). The plasmids were then sent for DNA sequencing (1st BASE, Singapore). The sequencing results were subsequently aligned with the digital template and analysed on Benchling platform.

pBbE6K (JBEI Part ID: JPUB 000054, *colE1* ori, *Kan*^*r*^), pBbE8K (JBEI Part ID: JPUB 000036, *colE1* ori, *Kan*^*r*^), pBbA6C (JBEI Part ID: JPUB 000056, *p15A* ori, *Cm*^*r*^), pBbA8C (JBEI Part ID: JPUB 000038, *p15A* ori, *Cm*^*r*^) and pNO4, which was a gift from Jeffrey Tabor (*pSC101 ori, Amp*^*r*^) (Addgene plasmid # 101066 ; http://n2t.net/addgene:101066 ; RRID:Addgene_101066), were used as the backbones in the plasmid constructions whenever necessary. Constitutive promoters J23101 (BBa_J23101) and gabDP2 (BBa_K3252022), pT7 promoter (BBa_R0085), double terminator 15T (BBa_B0015), rrnBT1 terminator (BBa_B0010), T7 terminator (BBa_K731721), ribosome binding site (rbs34) (BBa_B0034), green fluorescence protein gene *GFPmut3b* (BBa_E0040), red fluorescence protein gene *DsRed* (BBa_K2782004), sugar transport related sRNA gene (*SgrS)* (BBa_K581005) and the T7 RNA polymerase gene (BBa_I2032) were obtained from iGEM Registry of Standard Biological Parts (iGEM Foundation, Cambridge, MA, USA) (http://partsregistry.org) and used in plasmid constructions whenever necessary. The T7 RNA polymerase gene was split at the 563 (S) /564^th^ (E) location, to form the N-terminal T7 protein unit and C-terminal T7 protein unit, as guided by previous work when attempting to find the most optimum split sites (Baumschlager *et al*., 2017; Han *et al*., 2017). The GGSGG linker was obtained from a previous study (Baumschlager *et al*., 2017). The full length 371 amino acids (a.a) long *Tlpa36* gene (referred herein as *Tlpa)* and its pTlpa promoter were derived from the paper (Piraner *et al*., 2017), and synthesised as a gene fragment.

The plasmids P_GFP reporter_ was generated by inserting PCR amplified sequence (pT7-rbs34-GFP) into backbone pBbE6k. The plasmid P_gabDP2-NT7Tlpa-GFP_ was generated by inserting PCR amplified sequences (promoter gabDP2-rbsDefault-NT7-linker-Tlpa coil) into P_GFP reporter_ with multiple PCR steps. Similarly, Plasmids P_J23101-NT7Tlpa-GFP_ was generated by inserting PCR amplified sequences of promoter J23101. The plasmid P_J23101-CT7Tlpa_ was generated by inserting the PCR amplified sequence (J23101-rbs34-Tlpa coil-linker-CT7) into backbone pBbA8c. The plasmid P_J23101-rbs34-T7full_ was generated by inserting PCR amplified sequence (promoter J23101-rbs34-T7RNAP) into backbone pBbA8c. The plasmids P_J23101-NT7-GFP_ and P_J23101-CT7_ were generated by PCR-mediated excision of the Tlpa coiled-coil sequences from the original plasmids P_J23101-NT7Tlpa-GFP_ and P_J23101-CT7Tlpa_ respectively. The plasmids P_J23101-NT7-GFP_ and P_J23101-CT7_ were generated by PCR-mediated excision of the Tlpa coiled-coil sequences from the original plasmids P_J23101-NT7Tlpa-GFP_ and P_J23101-CT7Tlpa_ respectively. While, P_J23101-NT7Tlpa-J23101-CT7Tlpa_ was generated by inserting the PCR amplified sequence (J23101-rbs34-Tlpa coil-linker-CT7) originally from P_J23101-CT7Tlpa_ into backbone P_J23101-NT7Tlpa-GFP_. Each of the plasmids P_gabDP2-NT7Tlpa-GFP_ and P_J23101-NT7Tlpa-GFP_ was co-transformed with P_J23101-CT7Tlpa_ to form Thermal-T7RNAP(v1 and v2) systems. P_J23101-NT7Tlpa-J23101-CT7Tlpa_ was co-transformed with P_GFP reporter_ to form Thermal-T7RNAP(v3). P_J23101-rbs34-T7full_ was co-transformed with P_GFP reporter_ to form the full length T7-WT control. P_J23101-NT7Tlpa-GFP_ and P_J23101-CT7Tlpa_ was co-transformed to form the Split-T7 control. Next, the plasmids P_J23100-Tlpa_ and P_pTlpa-RFP_ were generated by inserting PCR amplified sequences (promoter J23100-rbs34-Tlpa36) and (promoter pTlpa-rbs34-RFP) into backbones pBbA8c and pBbE6k respectively. Correspondingly, the plasmids P_J23100-Tlpa_ and P_pTlpa-RFP_ were co-transformed to form the Tlpa system. In the growth inhibition experiments, the plasmid P_gabDP2-NT7Tlpa-pT7-sgrs-GFPreporter_ was generated by inserting PCR amplified sequence (rbs34-SgrS-15T-J23101-rbs34-GFP-rrnBT1) into backbone P_gabDP2-NT7Tlpa-GFP_ with multiple PCR steps. Its control plasmid P_gabDP2-NT7Tlpa-GFPreporter(control)_ was generated by PCR-mediated excision of the sequence (promoter pT7-rbs34-GFP-15T) from the backbone P_gabDP2-NT7Tlpa-GFP_ and the subsequent insertion of sequence (promoter J23101-rbs34-GFP-rrnBT1) into the same backbone. Both of the plasmids P_gabDP2-NT7Tlpa-pT7-sgrs-GFPreporter_ and P_gabDP2-NT7Tlpa-GFPreporter(control)_ were each co-transformed with P_J23101-CT7Tlpa_ in chemical competent *E*.*coli K-12 MG1655* cells to form the ThermalT7RNAP-SgrS system and the ThermalT7RNAP-SgrS (control) system respectively. Next, the plasmid P_pTlpa-sgrs-RFPreporter_ was generated by inserting PCR amplified sequence (SgrS-15T-J23101-rbs34) into backbone P_pTlpa-RFP_. Its control plasmid P_pTlpa-sgrs-RFPreporter(control)_ was generated by PCR-mediated excision of the promoter sequence pTlpa from the backbone P_pTlpa-RFP_ and the subsequent insertion of new promoter J23101 into the same backbone. Both of the plasmids P_pTlpa-sgrs-RFPreporter_ and P_pTlpa-sgrs-RFPreporter(control)_ were each co-transformed with P_J23100-Tlpa_ in *MG1655* cells to form the Tlpa-SgrS system and the Tlpa-SgrS (control) system respectively.

### Growth and characterisation

All chemicals were purchased from Sigma Aldrich (Sigma, MO, USA), unless stated otherwise. All glycerol stocks were prepared by mixing 500 μL overnight culture with 500 μL of sterilized 100% glycerol. Seed cultures from glycerol stocks were inoculated 20 hours overnight in 5 mL Invitrogen Luria Broth Base LB medium (ThermoFischer, USA) and supplemented with Kanamycin sulfate (Merck, Germany) (50 μg/mL), Chloramphenicol (25 μg/mL) and Ampicillin (100 μg/mL) whenever necessary and incubated in a mini NB-205 shaking incubator (BioTek, USA) at 40 °C, 225rpm.

To run the thermal regulation assay, 100 μL of overnight cultures were added to 5 mL fresh pre-warmed LB and grown for 90 mins at 40 °C at 225rpm. Subsequently, the refreshed cell cultures were dispensed into the wells of the 96-Well Non-Skirted PCR plate (Thermo Scientific, USA) at 100 μL. Final concentrations of 1 mM of IPTG (FirstBase, Singapore) and 0.2 % L(+) arabinose were added to some cell cultures which required chemical induction. Once 96-Well PCR plate had been loaded, the plate was sealed fully with adhesive film, and left to incubate inside the thermal cycler T100TM (Bio-Rad Laboratories, Hercules, CA, USA). Each row was programmed at the same temperature using the thermal ‘gradient’ function to create the desired temperature range from 30 to 42 °C. After the 18 hours incubation, the cell culture in each individual well was transferred into the 96-well microplate using a multi-channel pipette. The corresponding fluorescence intensities and optical density readings were read in the H1 Synergy (BioTek, USA) at the following settings: GFP gain: 75, GFP: excitation 485 nm, emission 528 nm; OD: 600nm.

For continuous kinetic experiment, the refreshed cell cultures were dispensed into the 96-well microplate at 300 μL each. The plate was covered with the plate lid and time series optical density and fluorescence were obtained at an interval of 10 minutes for a total duration of 12 hours and configured at double orbital shaking speed of 282 rpm continuously. Depending on the need of the studies, the temperature profiles were configured differently (at a fixed temperature of 30 °C /37 °C or transitioning between the two temperatures at intervals of 2/3 hours) in the microplate reader. Prior to the experiment, the system was pre-incubated at 40 °C to ensure tight repressibility.

### Automated colony screening for Thermal-T7RNAP mutants

Error-prone PCR was performed on the coiled-coil domains of the NT7 and CT7 protein fragments on plasmids P_gabDP2-NT7Tlpa-GFP_ and P_J23101-CT7Tlpa_ of the Thermal-T7RNAP(v1) system, using the GeneMorph II random mutagenesis kit (Agilent, USA). The generated PCR products were inserted back into the respective backbones using Gibson Assembly. The resultant mutant libraries were co-transformed with its un-mutated counterpart plasmid P_gabDP2-NT7Tlpa-GFP_ or P_J23101-CT7Tlpa_ respectively in *DH-10 Beta* cells in LB Agar. Approximately 10 rounds of mutagenesis were conducted to generate sufficient variants. After incubating overnight at 30 °C, each individual colony was isolated using the Rotor HDA colony picker (Singer instruments, UK). Using the colony picker, each colony was replicative plated onto three identical 384-format agar plates. These three identical agar plates were each grown overnight at 30 °C, 37 °C and 40 °C to screen for colonies with different activation and repressed temperatures. The next day, each identical colony under different temperatures was imaged in the colony picker upon illuminating with blue epifluorescence. Data processing was conducted to normalize the captured green fluorescence intensities with the corresponding size of individual colony (measured in mm^2^) and each colony was sorted and ranked by the respective fold changes in fluorescence intensities between 30°C/37°C and 30°C/40°C. Approximately 1700 colonies were screened in the library and 130 CT7 mutant colonies were shortlisted and quantified in the thermal regulation assay as described before. The thermal induction profiles generated were each fitted with a Hill’s equation and correspondingly the activity and performance are quantified with various ranking indexes such as fold change between 10% and 90% of fluorescence intensities, the steepness of transition and half-activation temperatures.

### Growth inhibition assay

Seed cultures of the GFP cells (ThermalT7RNAP-SgrS system) and the RFP cells (Tlpa-SgrS system), along with their control cells, were cultured 20 hours overnight in 5 ml of LB medium with the necessary antibiotics at their repressive temperature of 40 °C and 30 °C respectively. 50 μL of overnight cultures were added to 5 mL fresh pre-warmed M9 medium and grown for 90 mins at their repressive temperatures at 225 rpm. The M9 medium was prepared by adding 5x M9 salts, with final concentrations of 0.2% casamino acids, 100 μM CaCl_2_, 2 mM MgSO_4_ and 0.2% glucose into distilled water. The refreshed cell cultures were individually corrected to reach growth density of OD=0.1, before being dispensed in the 96-well microplate at their respective cell proportions into each well to form various co-cultures. The microplates were conducted in a continuous kinetic fashion at the required temperatures following the same microplate settings as described before.

### Computational modelling

A mechanistic model represented in the form of ordinary differential equations (ODEs) was formulated to describe the kinetics of the thermal-repressible split-T7RNAP fusion protein and used to examine the different gene circuit configurations (Supplementary Table S2-S4). The same model was also employed to gain insight into the dynamic behaviors of the 3-hour and 2-hour interval OFF-ON thermal duty cycles. To better quantify transcriptional kinetics, a simple model was adopted from Motta-Mena et al. (Motta-Mena *et al*., 2014) to derive the activation (*τ*_*ON*_) and deactivation (*τ*_*OFF*_) time constants from the protein synthesis rate profiles.

Further, growth models were developed based upon monoculture data to predict the combined growth profiles and individual growth profiles of the cell populations within the co-cultures, seeded at different initial cell proportions and temperatures (Supplementary Table S5-S6). As an extension, a phenomenological model was developed (Supplementary Table S7-S8) to correlate the cell growth with the reporting fluorescence.

To facilitate the model development process, several ‘modules’ (consisting of promoter-rbs-GFP) were characterised using microplate reader and their time-series profiles were fed into the BioModel Selection System (BMSS) (Yeoh et al., 2019) to identify an appropriate representative ODE model to be used for full model construction. These model parameters derived from the BMSS were estimated using a two-step optimization: differential evolution global optimizer followed by a constrained Nelder-Mead local optimizer (https://github.com/EngBioNUS/BMSSlib). The same optimization technique was employed for parameter inference in most of the developed models including the models used in predicting the co-culture dynamics.

To better capture the uncertainties of model parameter estimates, Bayesian parameter inference method was also deployed to infer the probability distributions of some parameters estimates as opposed to the singleton estimated parameters values (Supplementary Figure S8). This inference approach is closely relevance to the Bayesian interpretation of probability, in which it depends on prior knowledge or beliefs, evidence, and likelihood to infer the posterior distributions of the parameter’s estimates. The Metropolis-Hastings algorithm of the Markov chain Monte Carlo (MCMC) method was implemented in the parameterization process (Yildirim, 2012), which enables one to sample from distribution without having to compute all the high dimensional integrals that demands huge computational efforts. A normal distribution is assumed for all the priors with half of the individual mean values were assigned to the individual standard deviations. A detailed description of the Bayesian approach is included in Supplementary Note S2.

### Statistics

All data were shown as mean ± S.D (n=3). All samples were prepared in technical triplicates; Statistical significance was determined by performing a two-sample unpaired t-test using Microsoft Excel (Microsoft, USA), a prior F-test was conducted to reveal equal variance or unequal variance of the samples in comparison. Blanking was included in each experiment whereby the auto-fluorescence reading of the medium was recorded. The GFP/OD_600nm_ reading was calculated as fluorescence of (GFP_sample_ – GFP_blank_)/OD_600nm_ at each time point.

## REFERENCES

Abedi, M.H., Lee, J., Piraner, D.I., and Shapiro, M.G. (2020). Thermal Control of Engineered T-cells. ACS Synthetic Biology 9, 1941–1950. 10.1021/acssynbio.0c00238.

Aucoin, M.G., McMurray-Beaulieu, V., Poulin, F., Boivin, E.B., Chen, J., Ardelean, F.M., Cloutier, M., Choi, Y.J., Miguez, C.B., and Jolicoeur, M. (2006). Identifying conditions for inducible protein production in E. coli: combining a fed-batch and multiple induction approach. Microb Cell Fact 5, 27. 10.1186/1475-2859-5-27.

Baumschlager, A., Aoki, S.K., and Khammash, M. (2017). Dynamic Blue Light-Inducible T7 RNA Polymerases (Opto-T7RNAPs) for Precise Spatiotemporal Gene Expression Control. ACS Synth Biol 6, 2157–2167. 10.1021/acssynbio.7b00169.

Briand, L., Marcion, G., Kriznik, A., Heydel, J.M., Artur, Y., Garrido, C., Seigneuric, R., and Neiers, F. (2016). A self-inducible heterologous protein expression system in Escherichia coli. Sci Rep 6, 33037. 10.1038/srep33037.

Chait, R., Ruess, J., Bergmiller, T., Tkačik, G., and Guet, C.C. (2017). Shaping bacterial population behavior through computer-interfaced control of individual cells. Nature Communications 8, 1535. 10.1038/s41467-017-01683-1.

Chen, F., and Wegner, S.V. (2020). Blue-Light-Switchable Bacterial Cell-Cell Adhesions Enable the Control of Multicellular Bacterial Communities. ACS Synth Biol 9, 1169–1180. 10.1021/acssynbio.0c00054.

Dunstan, M.S., Robinson, C.J., Jervis, A.J., Yan, C., Carbonell, P., Hollywood, K.A., Currin, A., Swainston, N., Feuvre, R.L., Micklefield, J., et al. (2020). Engineering Escherichia coli towards de novo production of gatekeeper (2S)-flavanones: naringenin, pinocembrin, eriodictyol and homoeriodictyol. Synthetic Biology 5, 1–11. 10.1093/synbio/ysaa012.

Fang, Y., Wang, J., Ma, W., Yang, J., Zhang, H., Zhao, L., Chen, S., Zhang, S., Hu, X., Li, Y., and Wang, X. (2020). Rebalancing microbial carbon distribution for L-threonine maximization using a thermal switch system. Metabolic Engineering 61, 33–46. https://doi.org/10.1016/j.ymben.2020.01.009.

Fu, L., Gong, J., Gao, B., Ji, D., Han, X., and Zeng, L. (2020). Controlled expression of lysis gene E by a mutant of the promoter pL of the thermo-inducible λcI857-pL system. Journal of applied microbiology, 1–10.

Gal-Mor, O., Valdez, Y., and Finlay, B.B. (2006). The temperature-sensing protein TlpA is repressed by PhoP and dispensable for virulence of Salmonella enterica serovar Typhimurium in mice. Microbes Infect 8, 2154–2162. 10.1016/j.micinf.2006.04.015.

Gamboa, L., Phung, E.V., Li, H., Meyers, J.P., Hart, A.C., Miller, I.C., and Kwong, G.A. (2020). Heat-Triggered Remote Control of CRISPR-dCas9 for Tunable Transcriptional Modulation. ACS Chemical Biology 15, 533–542. 10.1021/acschembio.9b01005.

Goers, L., Freemont, P., and Polizzi, K.M. (2014). Co-culture systems and technologies: taking synthetic biology to the next level. J R Soc Interface 11, 1–13. 10.1098/rsif.2014.0065.

Greco, F.V., Grierson, C.S., and Gorochowski, T.E. (2020). Harnessing the central dogma for stringent multi-level control of gene expression. bioRxiv, 2020.2007.2004.187500. 10.1101/2020.07.04.187500.

Guarino, A., Fiore, D., Salzano, D., and di Bernardo, M. (2020). Balancing Cell Populations Endowed with a Synthetic Toggle Switch via Adaptive Pulsatile Feedback Control. ACS Synth Biol 9, 793–803. 10.1021/acssynbio.9b00464.

Han, T., Chen, Q., and Liu, H. (2017). Engineered Photoactivatable Genetic Switches Based on the Bacterium Phage T7 RNA Polymerase. ACS Synth Biol 6, 357–366. 10.1021/acssynbio.6b00248.

Harder, B.J., Bettenbrock, K., and Klamt, S. (2018). Temperature-dependent dynamic control of the TCA cycle increases volumetric productivity of itaconic acid production by Escherichia coli. Biotechnology and bioengineering 115, 156–164. 10.1002/bit.26446.

Hoynes-O’Connor, A., Hinman, K., Kirchner, L., and Moon, T.S. (2015). De novo design of heat-repressible RNA thermosensors in E. coli. Nucleic Acids Res 43, 6166–6179. 10.1093/nar/gkv499.

Hussaina, F., Guptab, C., Hirninga, A.J., Ottb, W., Matthewsa, K.S., Josi, K., and Bennett, M.R. (2014). Engineered temperature compensation in a synthetic genetic clock. PNAS 111, 972–977.

Keto-Timonen, R., Hietala, N., Palonen, E., Hakakorpi, A., Lindstrom, M., and Korkeala, H. (2016). Cold Shock Proteins: A Minireview with Special Emphasis on Csp-family of Enteropathogenic Yersinia. Front Microbiol 7, 1151. 10.3389/fmicb.2016.01151.

Kortmann, J., and Narberhaus, F. (2012). Bacterial RNA thermometers: molecular zippers and switches. Nature Reviews Microbiology 10, 255–265. 10.1038/nrmicro2730.

Krieger, A.G., Zhang, J., and Lin, X.N. (2021). Temperature regulation as a tool to program synthetic microbial community composition. Biotechnology and bioengineering 118, 1–12. https://doi.org/10.1002/bit.27662.

Liang, R., Liu, X., Liu, J., Ren, Q., Liang, P., Lin, Z., and Xie, X. (2007). A T7-expression system under temperature control could create temperature-sensitive phenotype of target gene in Escherichia coli J Microbiol Methods 68, 497–506. 10.1016/j.mimet.2006.10.016.

Liu, C.C., Jewett, M.C., Chin, J.W., and Voigt, C.A. (2018). Toward an orthogonal central dogma. Nat Chem Biol 14, 103–106. 10.1038/nchembio.2554.

Miller, I.C., Gamboa Castro, M., Maenza, J., Weis, J.P., and Kwong, G.A. (2018). Remote Control of Mammalian Cells with Heat-Triggered Gene Switches and Photothermal Pulse Trains. ACS Synth Biol 7, 1167–1173. 10.1021/acssynbio.7b00455.

Motta-Mena, L.B., Reade, A., Mallory, M.J., Glantz, S., Weiner, O.D., Lynch, K.W., and Gardner, K.H. (2014). An optogenetic gene expression system with rapid activation and deactivation kinetics. Nature Chemical Biology 10, 196–202. 10.1038/nchembio.1430.

Naik, R.R., Kirkpatrick, S.M., and Stone, M.O. (2001). The thermostability of an α-helical coiled-coil protein and its potential use in sensor applications. Biosensors and Bioelectronics 16, 1051–1057. https://doi.org/10.1016/S0956-5663(01)00226-3.

Naseri, G., and Koffas, M.A.G. (2020). Application of combinatorial optimization strategies in synthetic biology. Nature Communications 11, 2446. 10.1038/s41467-020-16175-y.

Negrete, A., Ng, W.I., and Shiloach, J. (2010). Glucose uptake regulation in E. coli by the small RNA SgrS: comparative analysis of E. coli K-12 (JM109 and MG1655) and E. coli B (BL21). Microb Cell Fact 9, 75. 10.1186/1475-2859-9-75.

Nikolic, N., Barner, T., and Ackermann, M. (2013). Analysis of fluorescent reporters indicates heterogeneity in glucose uptake and utilization in clonal bacterial populations. BMC Microbiology 13, 258. 10.1186/1471-2180-13-258.

Nistala, G.J., Wu, K., Rao, C.V., and Bhalerao, K.D. (2010). A modular positive feedback-based gene amplifier. Journal of Biological Engineering 4, 4. 10.1186/1754-1611-4-4.

Pearce, S.C., McWhinnie, R.L., and Nano, F.E. (2017). Synthetic temperature-inducible lethal gene circuits in Escherichia coli. Microbiology 163, 462–471. 10.1099/mic.0.000446.

Peng, Y.Y., Howell, L., Stoichevska, V., Werkmeister, J.A., Dumsday, G.J., and Ramshaw, J.A.M. (2012). Towards scalable production of a collagen-like protein from Streptococcus pyogenes for biomedical applications. Microb Cell Fact 11, 1–8.

Piraner, D.I., Abedi, M.H., Moser, B.A., Lee-Gosselin, A., and Shapiro, M.G. (2017). Tunable thermal bioswitches for in vivo control of microbial therapeutics. Nat Chem Biol 13, 75–80. 10.1038/nchembio.2233.

Piraner, D.I., Wu, Y., and Shapiro, M.G. (2019). Modular Thermal Control of Protein Dimerization. ACS Synthetic Biology 8, 2256–2262. 10.1021/acssynbio.9b00275.

Qing, G., Ma, L.C., Khorchid, A., Swapna, G.V., Mal, T.K., Takayama, M.M., Xia, B., Phadtare, S., Ke, H., Acton, T., et al. (2004). Cold-shock induced high-yield protein production in Escherichia coli. Nat Biotechnol 22, 877–882. 10.1038/nbt984.

Rodrigues, J.L., Couto, M.R., Araújo, R.G., Prather, K.L.J., Kluskens, L., and Rodrigues, L.R. (2017). Hydroxycinnamic acids and curcumin production in engineered Escherichia coli using heat shock promoters. Biochemical Engineering Journal 125, 41–49. https://doi.org/10.1016/j.bej.2017.05.015.

Rodrigues, J.L., and Rodrigues, L.R. (2018). Potential Applications of the Escherichia coli Heat Shock Response in Synthetic Biology. Trends Biotechnol 36, 186–198. 10.1016/j.tibtech.2017.10.014.

Roell, G.W., Zha, J., Carr, R.R., Koffas, M.A., Fong, S.S., and Tang, Y.J. (2019). Engineering microbial consortia by division of labor. Microbial Cell Factories 18, 35 10.1186/s12934-019-1083-3.

Salila Vijayalal Mohan, H.K., Chee, W.K., Li, Y., Nayak, S., Poh, C.L., and Thean, A.V.Y. (2020). A highly sensitive graphene oxide based label-free capacitive aptasensor for vanillin detection. Materials & Design 186, 108208. https://doi.org/10.1016/j.matdes.2019.108208.

Schramm, T., Lempp, M., Beuter, D., Sierra, S.G., Glatter, T., and Link, H. (2020). High-throughput enrichment of temperature-sensitive argininosuccinate synthetase for two-stage citrulline production in E. coli. Metabolic Engineering 60, 14–24. https://doi.org/10.1016/j.ymben.2020.03.004.

Segall-Shapiro, T.H., Meyer, A.J., Ellington, A.D., Sontag, E.D., and Voigt, C.A. (2014). A ‘resource allocator’ for transcription based on a highly fragmented T7 RNA polymerase. Mol Syst Biol 10, 742. 10.15252/msb.20145299.

Sen, S., Apurva, D., Satija, R., Siegal, D., and Murray, R.M. (2017). Design of a Toolbox of RNA Thermometers. ACS Synthetic Biology 6, 1461–1470. 10.1021/acssynbio.6b00301.

Shah, N.H., and Muir, T.W. (2014). Inteins: Nature’s Gift to Protein Chemists. Chem Sci 5, 446–461. 10.1039/C3SC52951G.

Singh, R., Kumar, M., Mittal, A., and Mehta, P.K. (2016). Microbial enzymes: industrial progress in 21st century. 3 Biotech 6, 174–174. 10.1007/s13205-016-0485-8.

Sorensen, H.P., and Mortensen, K.K. (2005). Soluble expression of recombinant proteins in the cytoplasm of Escherichia coli. Microb Cell Fact 4, 1. 10.1186/1475-2859-4-1.

Stephens, K., Pozo, M., Tsao, C.-Y., Hauk, P., and Bentley, W.E. (2019). Bacterial co-culture with cell signaling translator and growth controller modules for autonomously regulated culture composition. Nature Communications 10, 4129. 10.1038/s41467-019-12027-6.

Stirling, F., Bitzan, L., O’Keefe, S., Redfield, E., Oliver, J.W.K., Way, J., and Silver, P.A. (2017). Rational Design of Evolutionarily Stable Microbial Kill Switches. Mol Cell 68, 686–697 e683. 10.1016/j.molcel.2017.10.033.

Temme, K., Hill, R., Segall-Shapiro, T.H., Moser, F., and Voigt, C.A. (2012). Modular control of multiple pathways using engineered orthogonal T7 polymerases. Nucleic Acids Res 40, 8773–8781. 10.1093/nar/gks597.

Valdez-Cruz, N.A., Caspeta, L., Pérez, N.O., Ramírez, O.T., and Trujillo-Roldán, M.A. (2010). Production of recombinant proteins in E. coli by the heat inducible expression system based on the phage lambda pL and/or pR promoters. Microb Cell Fact 9, 1–16.

Wadler, C.S., and Vanderpool, C.K. (2007). A dual function for a bacterial small RNA: SgrS performs base pairing-dependent regulation and encodes a functional polypeptide. Proc Natl Acad Sci U S A 104, 20454–20459. 10.1073/pnas.0708102104.

Wang, W., Li, Y., Wang, Y., Shi, C., Li, C., Li, Q., and Linhardt, R.J. (2018). Bacteriophage T7 transcription system: an enabling tool in synthetic biology. Biotechnol Adv 36, 2129–2137. 10.1016/j.biotechadv.2018.10.001.

Wang, X., Han, J.-N., Zhang, X., Ma, Y.-Y., Lin, Y., Wang, H., Li, D.-J., Zheng, T.-R., Wu, F.-Q., Ye, J.-W., and Chen, G.-Q. (2021). Reversible thermal regulation for bifunctional dynamic control of gene expression in Escherichia coli. Nature Communications 12, 1411. 10.1038/s41467-021-21654-x.

Wei, Y., Murphy, E.R., Larramendy, M., and Soloneski, S. (2016). Temperature-dependent regulation of bacterial gene expression by RNA thermometers. Nucleic Acids––from Basic Aspects to Laboratory Tools Specific. IntechOpen, London, 157–181.

Xu, S., Wang, Q., Zeng, W., Li, Y., Shi, G., and Zhou, J. (2020). Construction of a heat-inducible Escherichia coli strain for efficient de novo biosynthesis of l-tyrosine. Process Biochemistry 92, 85–92. https://doi.org/10.1016/j.procbio.2020.02.023.

Yang Zheng, Fankang Meng, Zihui Zhu, Weijia Wei, Zhi Sun, Jinchun Chen, Bo Yu, Chunbo Lou, and Chen, G.-Q. (2019). A tight cold-induicble switch by coupling thermosensitive transcriptional and proteolytic regulatory parts. Nucleic Acids Research 47, e137. 10.1093/nar/gkz785.

Yeoh, J.W., Ng, K.B.I., Teh, A.Y., Zhang, J., Chee, W.K.D., and Poh, C.L. (2019). An Automated Biomodel Selection System (BMSS) for Gene Circuit Designs. ACS Synthetic Biology 8, 1484–1497. 10.1021/acssynbio.8b00523.

Yildirim, I. (2012). Bayesian inference: Metropolis-hastings sampling (Dept. of Brain and Cognitive Sciences, Univ. of Rochester, Rochester, NY).

Zhang, C., Seow, V.Y., Chen, X., and Too, H.-P. (2018). Multidimensional heuristic process for highyield production of astaxanthin and fragrance molecules in Escherichia coli. Nature Communications 9, 1858. 10.1038/s41467-018-04211-x.

Zhao, E.M., Suek, N., Wilson, M.Z., Dine, E., Pannucci, N.L., Gitai, Z., Avalos, J.L., and Toettcher, J.E. (2019). Light-based control of metabolic flux through assembly of synthetic organelles. Nat Chem Biol 15, 589–597. 10.1038/s41589-019-0284-8.

Zhao, E.M., Zhang, Y., Mehl, J., Park, H., Lalwani, M.A., Toettcher, J.E., and Avalos, J.L. (2018). Optogenetic regulation of engineered cellular metabolism for microbial chemical production. Nature 555, 683–687. 10.1038/nature26141.

