## Supplementary Information for "Highly Reversible Tunable Thermal-repressible Split-T7 RNA polymerases (Thermal-T7RNAPs) for dynamic gene regulation"

### SUPPLEMENTARY TABLES

**Supplementary Table S1.** The temperature dose-response behaviours of different genetic systems represented by Hill equations[a].

| Name of system | $n$ | $K_m$ | $K_{inh}$ |
|---|---|---|---|
| Full length T7-WT | 1.8990e+02 | 4.0846e+01 | 5.6382e-01 |
| Split-T7 (no Tlpa coil) | 5.4240e+01 | 3.7661e+01 | 7.5484e-01 |
| Thermal-T7RNAP(v1) | 6.9398e+01 | 3.7527e+01 | 9.5735e-01 |
| Thermal-T7RNAP(v2) | 1.2297e+02 | 3.8368e+01 | 4.2718e-01 |
| Thermal-T7RNAP(v3) | 3.5636e+01 | 3.6340e+01 | 9.7089e-01 |

[a]The Hill equation is given as $\left(1 - K_{inh}\left(\frac{T^n}{T^n+{K_m}^n}\right)\right)$, where $K_{inh}$ indicates the maximum repression capacity; $K_m$ and $n$ represent the half-activation temperature and the hill coefficient respectively.

**Supplementary Table S2.** The model formulation used to elucidate the kinetics of the thermal-repressible Split-T7RNAP protein fusion is governed by a cascade of ODEs as given below:

| Mathematical Equations | No. |
|---|---|
| $\frac{d[mRNA]_{NT7}}{dt} = \alpha_{mRNANT7} - \gamma_{mRNA}[mRNA]_{NT7}$ | Eq. S1 |
| $\frac{d[Pep]_{NT7}}{dt} = \alpha_{PepNT7}[mRNA]_{NT7} - K_{mat}[Pep]_{NT7}$ | Eq. S2 |
| $\frac{d[NT7]}{dt} = K_{mat}[Pep]_{NT7} - \gamma_{Pep}[NT7] - K_b(1 - K_{inh}(state))[NT7][CT7] + K_{ub}[T7]$ | Eq. S3 |
| $\frac{d[mRNA]_{CT7}}{dt} = \alpha_{mRNACT7} - \gamma_{mRNA}[mRNA]_{CT7}$ | Eq. S4 |
| $\frac{d[Pep]_{CT7}}{dt} = \alpha_{PepCT7}[mRNA]_{CT7} - K_{mat}[Pep]_{CT7}$ | Eq. S5 |
| $\frac{d[CT7]}{dt} = K_{mat}[Pep]_{CT7} - \gamma_{Pep}[CT7] - K_b(1 - K_{inh}(state))[NT7][CT7] + K_{ub}[T7]$ | Eq. S6 |
| $\frac{d[T7]}{dt} = K_b(1 - K_{inh}(state))[NT7][CT7] - K_{ub}[T7]$ | Eq. S7 |
| $\frac{dmRNA_{GFP}}{dt} = \alpha_{mRNAGFP}\left(\frac{[T7]}{[T7] + K_m}\right) - \gamma_{mRNA}[mRNA]_{GFP}$ | Eq. S8 |
| $\frac{d[GFP]}{dt} = \alpha_{PepGFP}[mRNA]_{GFP} - \gamma_{Pep}[GFP]$ | Eq. S9 |

where $\alpha_{mRNAX}$ and $\alpha_{PepX}$ for X = $NT7$, $CT7$, $GFP$ represent the promoter-dependent mRNA synthesis rate and ribosome binding site (RBS)-dependent protein translation rate for the individual split N-terminal and C-terminal T7 proteins fragments, and the T7-driven green fluorescent protein GFP, respectively. $\gamma_{mRNA}$ and $\gamma_{Pep}$ denote the mRNA and protein degradation rates, whereas $K_{mat}$ refers to the protein maturation time. $K_b$ and $K_{ub}$ indicate the binding and unbinding rate constants for the fusion reaction of the Split-T7 proteins fragments, and $K_m$ refers to the half activation T7RNAP concentration. The $state$ represents the temperature control where 1 is used to denote 37 °C, and 0 indicates 30 °C, and $K_{inh}$ refers to the maximal repression capacity from 0 to 1 induced by the high temperature of 37 °C.

**Supplementary Table S3.** Parameters derived from the kinetic model presented in the previous supplementary table.

| Parameters | Continuous | 3hr (OFF-ON) |
|---|---|---|
| $\alpha_{mRNANT7}$ | 7.0543e-6 | Maintained |
| $\gamma_{mRNA}$ | 1.386e-01 | Maintained |
| $\alpha_{PepNT7}$ | 1.3213e-02 | Maintained |
| $\gamma_{Pep}$ | 7.18e-03 | Maintained |
| $K_{mat}$ | 4.68e-04 | Maintained |
| $\alpha_{mRNACT7}$ | 7.0543e-6 | Maintained |
| $\alpha_{PepCT7}$ | 1.01e-02 | Maintained |
| $K_b$ | 3.123e+02 | 1.0042e+4 |
| $K_{ub}$ | 6.097e-03 | 4.81e-01 |
| $K_b/K_{ub}$ | 5.1147e+04 | 2.0877e+04 |
| $K_{inh}$ | 6.05e-01 | 6.50e-01 |
| $\alpha_{mRNAGFP}$ | 1.833e-05 | 2.317e-05 |
| $K_m$ | 1e-06 | Maintained |
| $\alpha_{PepGFP}$ | 1.01e-02 | Maintained |

**Supplementary Table S4.** Parameters used in simulating the different gene circuit configurations.

| Circuit Configurations | Parameters | Values |
|---|---|---|
| Thermal-T7RNAP(v1) | $\alpha_{mRNANT7}$ | 9.829e-6 x 0.1357 |
| | $\alpha_{PepNT7}$ | 0.0101 x 1.3082 |
| | $\alpha_{mRNACT7}$ | 9.829e-6 x 0.7177 |
| | $\alpha_{PepCT7}$ | 0.0101 |
| Thermal-T7RNAP(v2) | $\alpha_{mRNANT7}$ | 9.829e-6 |
| | $\alpha_{PepNT7}$ | 0.0101 x 1.3082 |
| | $\alpha_{mRNACT7}$ | 9.829e-6 x 0.7177 |
| | $\alpha_{PepCT7}$ | 0.0101 |
| Thermal-T7RNAP(v3) | $\alpha_{mRNANT7}$ | 9.829e-6 x 0.7177 |
| | $\alpha_{PepNT7}$ | 0.0101 x 1.3082 |
| | $\alpha_{mRNACT7}$ | 9.829e-6 x 0.7177 |
| | $\alpha_{PepCT7}$ | 0.0101 |

To better account for the effect of gene length on mRNA and protein synthesis rates, all mRNA synthesis rates for NT7, $\alpha_{mRNANT7}$ were multiplied by a coefficient of 0.9663 whereas the corresponding protein synthesis rates for NT7 were multiplied by a coefficient of 0.9346. These coefficients were estimated using the integrative host-circuit model (1) taken into consideration of the longer amino acid length of NT7-Tlpa coil proteins (848 aa) and the shorter length of CT7-Tlpa coil proteins (604 aa). To predict the system behaviours at 42 °C, the repressed terms, $K_{inh}(state)$ from Eq. S3, S6, and S7 were replaced by the Thermal-T7RNAP(v3) dose-response equation's parameters: $K_m$ and $n$ , as derived from the data in the thermal regulation assay.

**Supplementary Table S5.** Mathematical Formulations describing the co-culture growth dynamics for cells carrying ThermalT7RNAP-SgrS system and Tlpa-SgrS system respectively, and for the control cells without SgrS expression.

| Conditions | Mathematical Equations | No. |
|---|---|---|
| Co-culture control cells (no SgrS) | $\frac{dOD_{600_{T7}}}{dt} = \mu_{maxT7}\left(\frac{T^n}{T^n + K_{mu}{}^n}\right)\left(\frac{[Nutr]}{[Nutr] + K_n}\right)OD_{600_{T7}}$ | Eq. S10 |
| | $\frac{dOD_{600_{TLPA}}}{dt} = \mu_{maxTLPA}\left(\frac{T^n}{T^n + K_{mu}{}^n}\right)\left(\frac{[Nutr]}{[Nutr] + K_n}\right)OD_{600_{TLPA}}$ | Eq. S11 |
| | $\frac{d[Nutr]}{dt} = -\frac{1}{Y_{Nutr}}dOD_{600_{T7}}C_{OD} - \frac{1}{Y_{Nutr}}dOD_{600_{TLPA}}C_{OD}$ | Eq. S12 |
| Co-culture cells (with SgrS) | $[sgrS]_{T7} = \left(\frac{K_{mtT7}{}^{nT7}}{K_{mtT7}{}^{nT7} + T^{nT7}}\right)\left(\frac{1}{1 + e^{(K_{mT7} - t)}}\right)$ | Eq. S13 |
| | $K_{nT7} = K_n + K_{inhT7}\left(\frac{[sgrS]_{T7}}{[sgrS]_{T7} + K_{sT7}}\right)$ | Eq. S14 |
| | $\frac{dOD_{600_{T7}}}{dt} = \mu_{maxT7}\left(\frac{T^n}{T^n + K_{mu}{}^n}\right)\left(\frac{[Nutr]}{[Nutr] + K_{nT7}}\right)OD_{600_{T7}}$ | Eq. S15 |
| | $[sgrS]_{TLPA} = \left(\frac{T^{nTLPA}}{K_{mtTLPA}{}^{nTLPA} + T^{nTLPA}}\right)\left(\frac{1}{1 + e^{(K_{mTLPA} - t)}}\right)$ | Eq. S16 |
| | $K_{nTLPA} = K_n + K_{inhTLPA}\left(\frac{[sgrS]_{TLPA}}{[sgrS]_{TLPA} + K_{sTLPA}}\right)$ | Eq. S17 |
| | $\frac{dOD_{600_{TLPA}}}{dt} = \mu_{maxTLPA}\left(\frac{T^n}{T^n + K_{mu}{}^n}\right)\left(\frac{[Nutr]}{[Nutr] + K_{nTLPA}}\right)OD_{600_{TLPA}}$ | Eq. S18 |
| | $\frac{d[Nutr]}{dt} = -\frac{1}{Y_{Nutr}}dOD_{600_{T7}}C_{OD} - \frac{1}{Y_{Nutr}}dOD_{600_{TLPA}}C_{OD}$ | Eq. S19 |

where $OD_{600_{T7}}$ and $OD_{600_{TLPA}}$ are the OD growth for cells carrying ThermalT7RNAP-SgrS system and Tlpa-SgrS system respectively; $\mu_{maxT7}$ and $\mu_{maxTLPA}$ are their respective maximum specific growth rates; $K_{mu}$ and $n$ represent the half-activation parameter and the slope of the default temperature-sensitive growth rate response; $Nutr$, $K_n$, $K_{nT7}$, and $K_{nTLPA}$ represent nutrient which is glucose in this case, the nutrient-sensitive half-activation parameters for control, and after considering SgrS effects on glucose uptake for T7, and Tlpa cells respectively; $Y_{Nutr}$ is the yield coefficient, $C_{OD}$ is the conversion factor from OD to g/L; $K_{mtT7}$, $nT7$, $K_{mtTLPA}$, and $nTLPA$ are the half-activation coefficients and slopes describing the temperature-dependent SgrS expressions for the two systems; Boltzmann function was used to feature the expression dynamics of SgrS, where $K_{mT7}$ and $K_{mTLPA}$ denote the time required for half-activated SgrS expressions; $K_{inhT7}$ and $K_{inhTLPA}$ indicate the maximum shift in glucose uptake sensitivity due to SgrS; whereas $K_{sT7}$ and $K_{sTLPA}$ refer to the half-activation effect of SgrS on shifting the sensitivity.

**Supplementary Table S6.** Model parameters of the formulations given in the previous supplementary table for the first dataset and after model adjustment, for the second dataset which included the mutant systems.

| Parameters | Dataset 1 | Dataset 2 | Units |
|---|---|---|---|
| $\mu_{maxT7}$ | 1.6149e-02 | Maintained | min-1 |
| $K_{mu}$ | 2.9238e+01 | 2.8212e+01 | °C |
| $n$ | 1.0916e+01 | 7.8161 | - |
| $K_n$ | 7.5869e-01 | Maintained | gL-1 |
| $\mu_{maxTLPA}$ | 1.3739e-02 | 1.4256e-2,<br>1.3085e-2 (control) | min-1 |
| $Y_{Nutr}$ | 1.2184e-01 | Maintained | - |
| $C_{OD}$ | 0.37 | Maintained | gL-1 |
| $K_{mtT7}$ | 30 | 3.7738e+01 (Original)<br>3.3004e+01 (Mut1)<br>3.3287e+01 (Mut2) | °C |
| $nT7$ | 4.3296 | 1.1472e+02 (Original)<br>1.3711e+02 (Mut1)<br>1.3272e+02 (Mut2) | - |
| $K_{mT7}$ | 8.82287e+01 | 1.3595e+01 | min |
| $K_{inhT7}$ | 5.2284 | 5.7007e-01 | gL-1 |
| $K_{sT7}$ | 1 | 4.2221e-07 | - |
| $K_{mtTLPA}$ | 3.9999e+01 | N/A | °C |
| $nTLPA$ | 1.4997e+02 | N/A | - |
| $K_{mTLPA}$ | 1.8447e+02 | N/A | min |
| $K_{inhTLPA}$ | 6.2513 | N/A | gL-1 |
| $K_{sTLPA}$ | 0.8857 | N/A | - |

**Supplementary Table S7.** Mathematical Formulations describing the correlations between the $OD_{600}$ growth and measured green or red fluorescence levels for respective monocultures at three different temperatures (30 °C, 37 °C, 40 °C).

| Conditions | Mathematical Equations | No. |
|---|---|---|
| GFP control and RFP control cells (No SgrS) | $K_m = K_{m0} - K_{inh}\left(\frac{T^{n1}}{T^{n1} + {K_{mt}}^{n1}}\right)$ | Eq. S20 |
| | $n = n_0 + n_{inh}\left(\frac{{K_{mt}}^{n1}}{{K_{mt}}^{n1} + T^{n1}}\right)$ | Eq. S21 |
| | $OD_{600} = \frac{OD_{max}}{1 + e^{\left(\frac{K_m - FP}{n}\right)}}$ | Eq. S22 |
| For GFP cell (with SgrS) | $K_m = K_{m0} - K_{inh}\left(\frac{T^{n1}}{T^{n1} + {K_{mt}}^{n1}}\right) - K_{inh2}\left(\frac{{K_{mt2}}^{n2}}{T^{n2} + {K_{mt2}}^{n2}}\right)$ | Eq. S23 |
| | $n = n_0 + n_{inh}\left(\frac{{K_{mt}}^{n1}}{{K_{mt}}^{n1} + T^{n1}}\right) + n_{inh3}\left(\frac{T^{n3}}{{K_{mt3}}^{n3} + T^{n3}}\right)$ | Eq. S24 |
| | $OD_{600} = \frac{OD_{max}}{1 + e^{\left(\frac{K_m - FP}{n}\right)}}$ | Eq. S25 |
| For RFP cell (with SgrS) | $K_m = K_{m0} - K_{inh}\left(\frac{T^{n1}}{T^{n1} + {K_{mt}}^{n1}}\right) - K_{inh2}\left(\frac{{K_{mt2}}^{n2}}{T^{n2} + {K_{mt2}}^{n2}}\right)$ | Eq. S26 |
| | $n = n_0 + n_{inh}\left(\frac{{K_{mt}}^{n}}{{K_{mt}}^{n} + T^{n}}\right) - n_{inh3}\left(\frac{{K_{mt3}}^{n3}}{{K_{mt3}}^{n3} + T^{n3}}\right) + n_{inh4}\left(\frac{T^{n4}}{{K_{mt4}}^{n4} + T^{n4}}\right)$ | Eq. S27 |
| | $OD_{600} = \frac{OD_{max}}{1 + e^{\left(\frac{K_m - FP}{n}\right)}}$ | Eq. S28 |

Here, the $OD_{600}$ was presented as a function of fluorescence level expressed constitutively by a Boltzmann function, where $OD_{max}$ is the maximum achievable OD, $K_m$ and $n$ are the corresponding half activation coefficient and slope, whereas $K_{m0}$ and $n_0$ refer to their basal values; $K_{inh}$ and $K_{inh2}$ indicate the left shift in $K_m$ due to increase and decrease in temperature respectively; $n_{inh}$, $n_{inh3}$, $n_{inh4}$ refer to the change in slope cause by temperature; and $K_{mt}$, $K_{mt2}$, $K_{mt3}$, $K_{mt4}$, $n1$, $n2$, $n3$, $n4$ are the corresponding temperature-sensitive half-activation or inhibition coefficients and their slopes.

**Supplementary Table S8.** Parameters derived from the correlations between the $OD_{600}$ growth and measured green or red fluorescence level as derived from the previous supplementary table.

| Parameters | GFP cells (control) | RFP cells (control) | GFP cells | RFP cells | Units |
|---|---|---|---|---|---|
| $K_{m0}$ | 2.905e+04 | 1.118e+03 | maintained | maintained | a.u. |
| $K_{inh}$ | 1.888e+04 | 4.924e+02 | maintained | maintained | a.u. |
| $K_{mt}$ | 3.068e+01 | 3.591e+01 | maintained | maintained | °C |
| $n1$ | 5.824e+00 | 7.690e+01 | maintained | maintained | - |
| $n_0$ | 4.555e+03 | 1.435e+02 | maintained | maintained | a.u. |
| $n_{inh}$ | 2.132e+03 | 2.632e+02 | maintained | maintained | a.u. |
| $OD_{max}$ | 7.4325e-01 | 7.685e-01 | maintained | maintained | a.u. |
| $K_{inh2}$ | N/A | N/A | 4.602e+03 | 4.706e+02 | a.u. |
| $K_{mt2}$ | N/A | N/A | 3.048e+01 | 3.601e+01 | °C |
| $n2$ | N/A | N/A | 9.313e+01 | 2.287e+01 | - |
| $n_{inh3}$ | N/A | N/A | 1.290e+03 | 2.267e+02 | a.u. |
| $K_{mt3}$ | N/A | N/A | 3.109e+01 | 3.275e+01 | °C |
| $n3$ | N/A | N/A | 9.999e+01 | 1.807e+01 | - |
| $n_{inh4}$ | N/A | N/A | N/A | 9.258e+02 | a.u. |
| $K_{mt4}$ | N/A | N/A | N/A | 4.0e+01 | °C |
| $n4$ | N/A | N/A | N/A | 4.477e+01 | - |

### SUPPLEMENTARY FIGURES

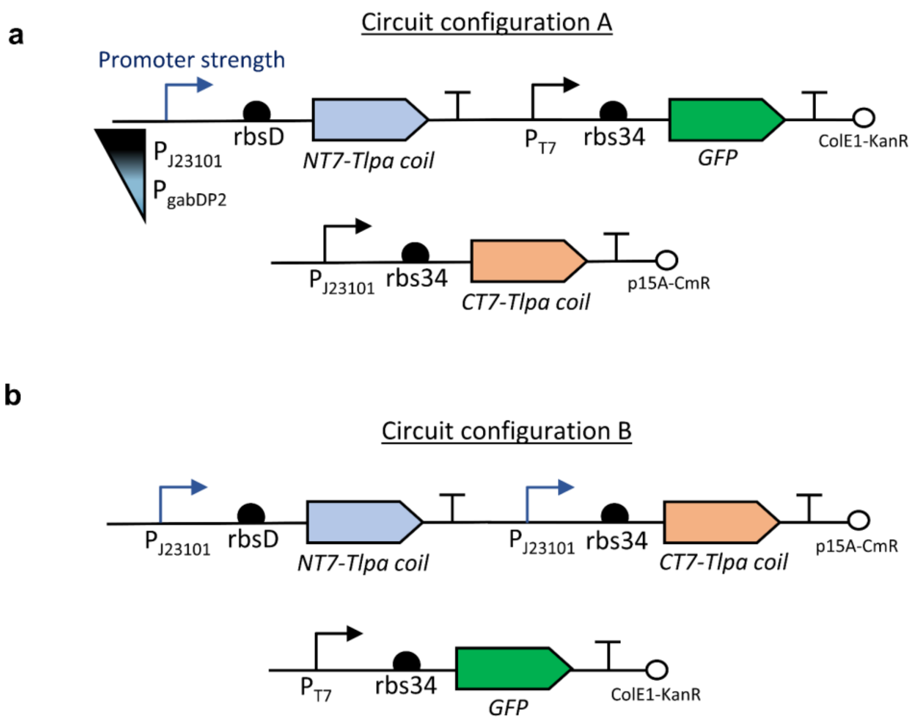


**Supplementary Figure S1.** Genetic circuits of Thermal-T7RNAP systems. (a) Gene configuration A. In Thermal-T7RNAP(v1) and Thermal-T7RNAP(v2) respectively, constitutive promoters of varying strengths (gabDP2-'low' and J23101-'high') are used in control of the expression levels of NT7-Tlpa coil protein, while the CT7-Tlpa coil protein is being expressed at a constant level using J23101 promoter on another plasmid of similar copy number. (b) Gene configuration B. In Thermal-T7RNAP(v3), the NT7-Tlpa coil and CT7-Tlpa coil are expressed on the same plasmid under J23101 promoters.

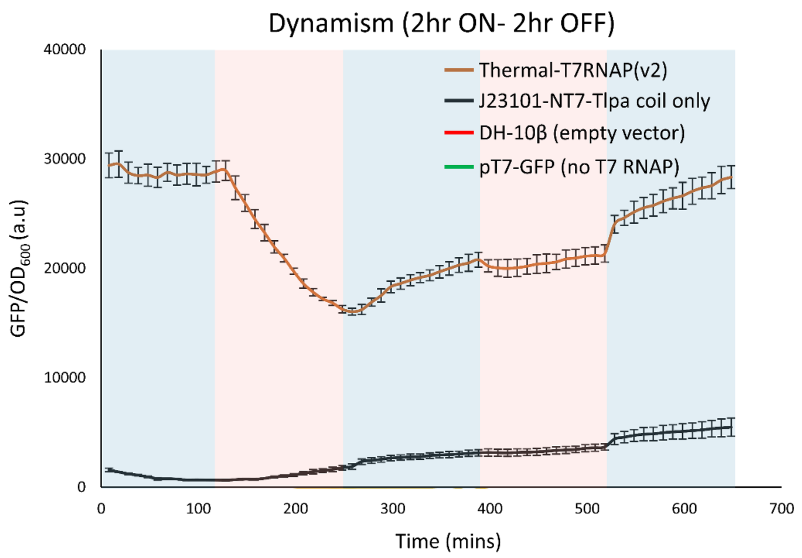


**Supplementary Figure S2.** Fluorescence expressions oscillating between 30 °C and 37 °C at 2hr-interval. Thermal-T7RNAP(v2), along with various controls such as the J23101-NT7-Tlpa coil system which allows us to study GFP expressions in the absence of the CT7 counterpart, the pT7-GFP system which does not contain T7RNAP expression and the background fluorescence emitted from the empty vector (DH-10β). The latter controls did not produce observable GFP expressions. Data information: All data are represented as mean ± S.D. (n = 3).

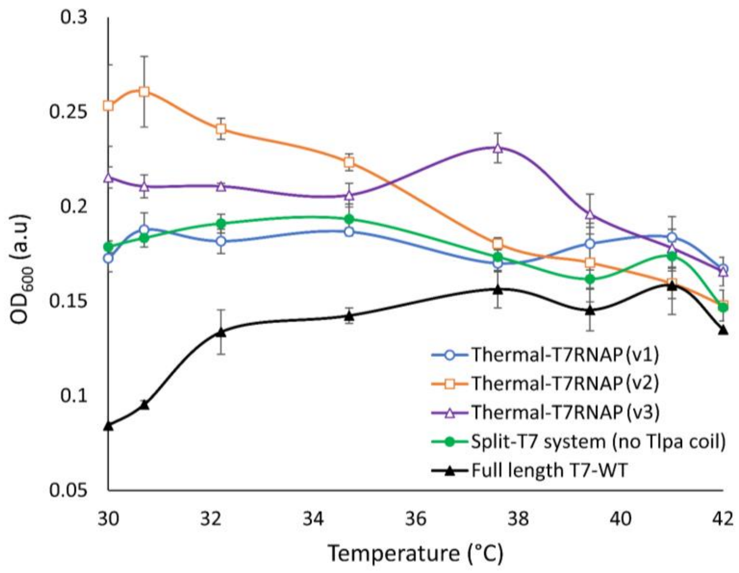


**Supplementary Figure S3.** Steady state growth values (after 18 hours) for the various Thermal-T7RNAP systems at different temperatures as performed in the thermal regulation assay. Data information: All data are represented as mean ± S.D. (n = 3).

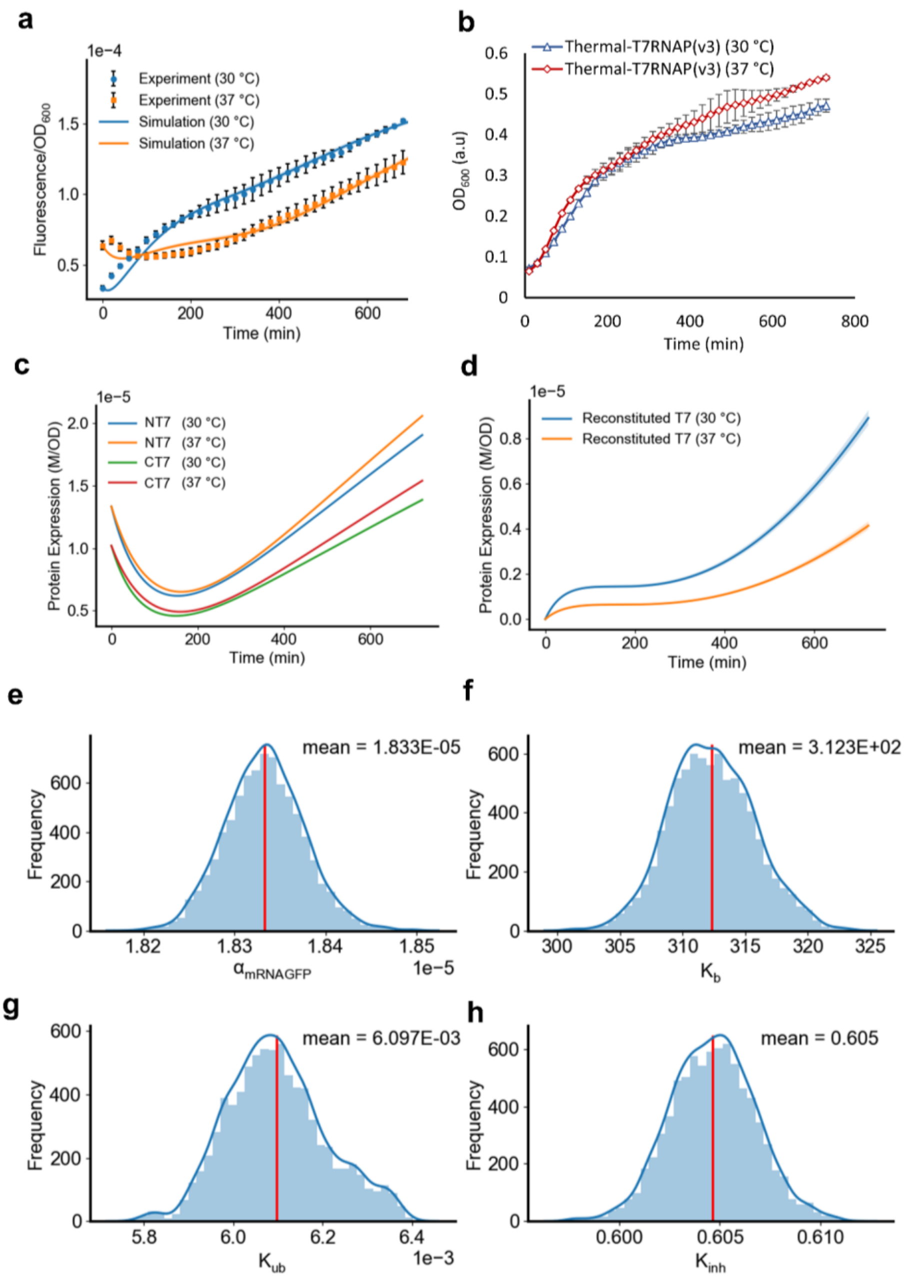


**Supplementary Figure S4.** (a) Model simulations of Thermal-T7RNAP(v3) to describe the continuous data characterized at 30 °C and 37 °C across a 12-hour timespan in comparison with experimental recordings. (b) Corresponding growth profile at different temperatures. (c) The predicted unbounded NT7 and CT7 proteins and (d) the reconstituted T7RNAP protein expression profiles. (e-h) The posterior distributions of the parameter estimates obtained from the Bayesian parameter inference method. Data information: Experimental data are represented as mean ± S.D. (n = 3).

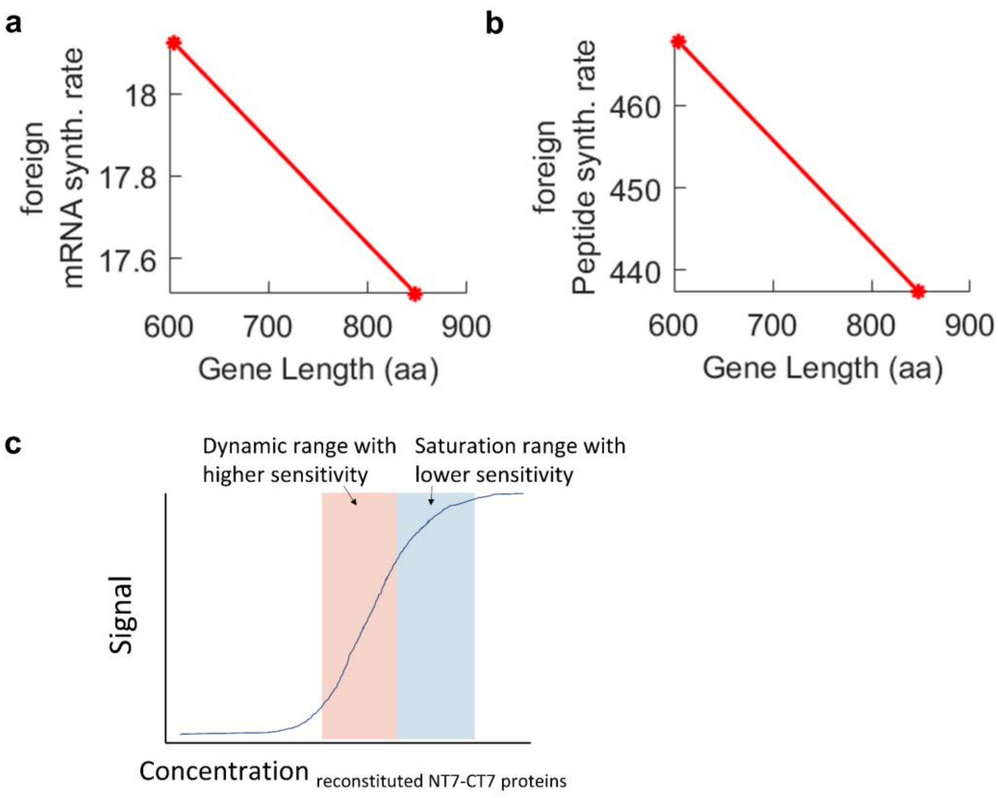


**Supplementary Figure S5.** The impacts of gene length represented as amino acid length (aa) on the (a) mRNA and (b) protein synthesis rates of the heterologous expression of the individual T7 protein fragments. The predictions were estimated using the integrative host-circuit model (1). (c) Sensitivity curve of NT7 and CT7 protein fragments which are bounded together to form an active polymerase.

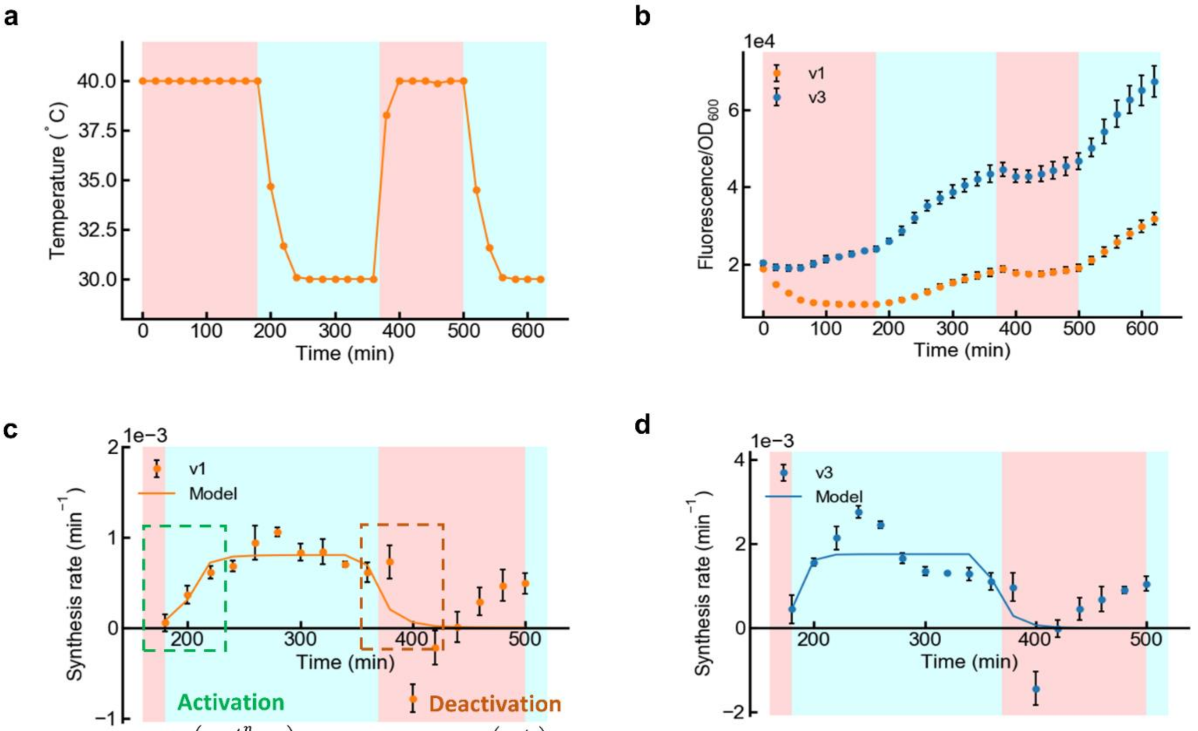


$$Y_{max}\left(\frac{t^n}{t^n+\tau_{ON}{}^n}\right) \qquad Y_{max}\exp\left(\frac{-t}{\tau_{OFF}}\right)$$

**Supplementary Figure S6.** (a) Measured temperature profile when alternating between 40 °C and 30 °C, over a 10-hour period. (b) Absolute Fluorescence expression profiles for Thermal-T7RNAP(v1) and Thermal-T7RNAP(v3). Derivation of activation and deactivation time constants ($\tau_{ON}$ and $\tau_{OFF}$) for (c) Thermal-T7RNAP(v1) and (d) Thermal-T7RNAP(v3) based on the synthesis rate profiles under the described temperature profile. In the equations, $\gamma_{max}$ refers to the maximum protein synthesis rate and n denotes the Hill coefficient (n = 4). Data information: All data are represented as mean ± S.D. (n = 3).

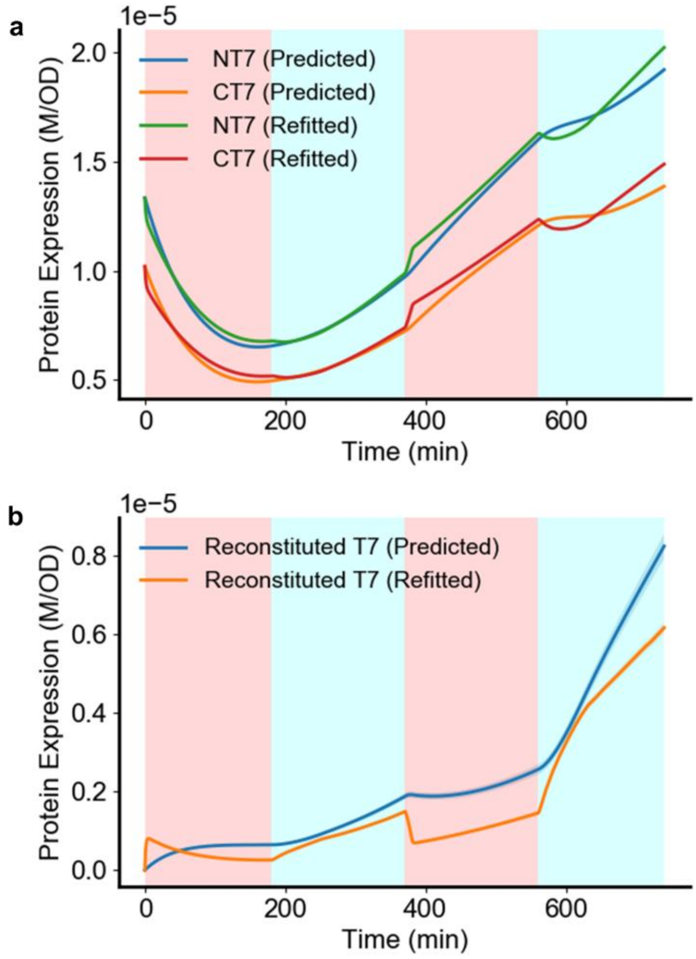


**Supplementary Figure S7.** The model predicted expression profiles for the split protein fragments and the reconstituted T7RNAP under the 3hr-OFF-ON-OFF-ON simulation dynamics. (a) The expression profiles of the unbounded NT7 and CT7 protein fragments. The predicted profiles were inferred from the continuous data whilst the refitted profiles were obtained from the refitted model that well captured the expression dynamics. (b) The predicted and the refitted model simulations for the reconstituted T7 in 3hr interval OFF-ON dynamics.

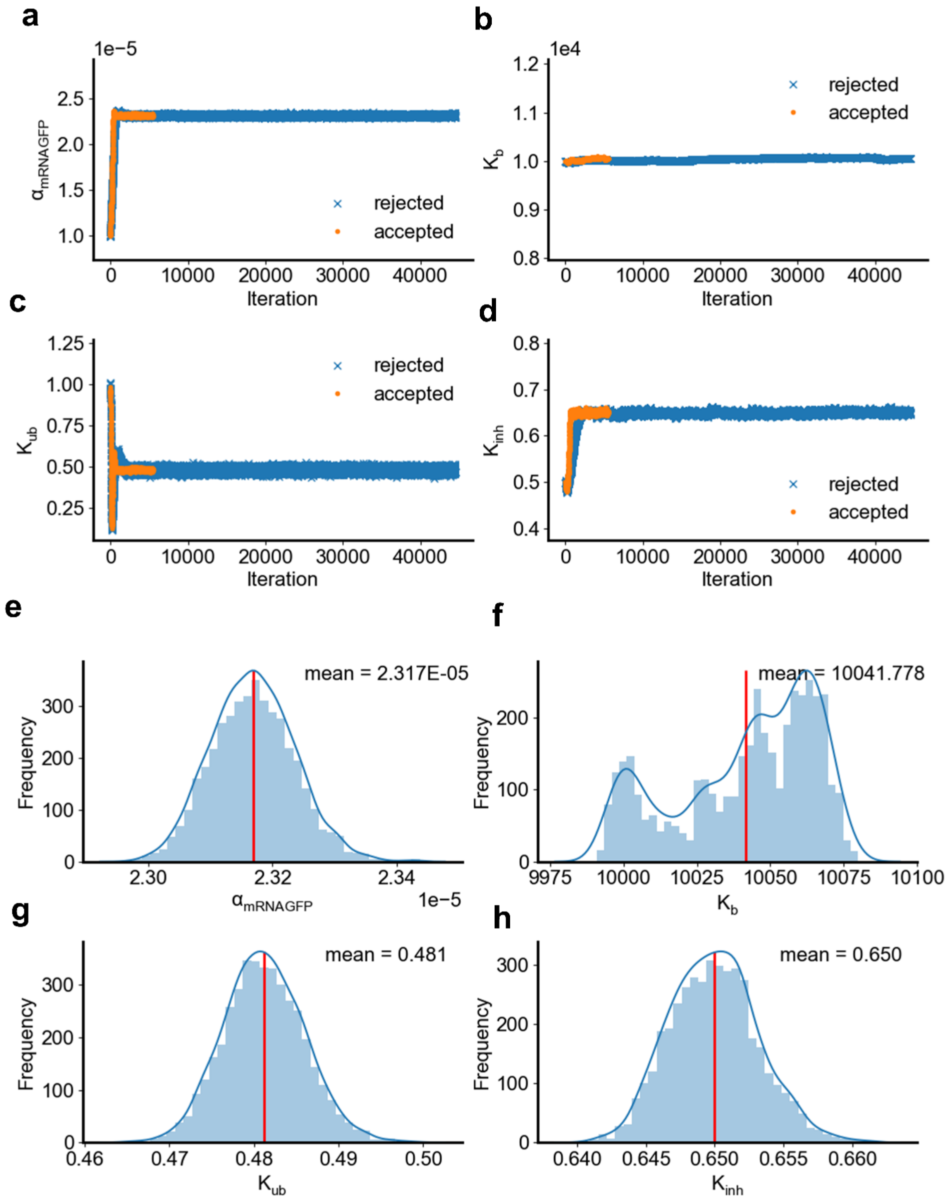


**Supplementary Figure S8.** (a-d) Trace plots for the inferred parameters obtained from the Bayesian parameter inference method. (e-h) Posterior plots for the parameters estimates after refitting the 3hr-OFF-ON dynamic profile.

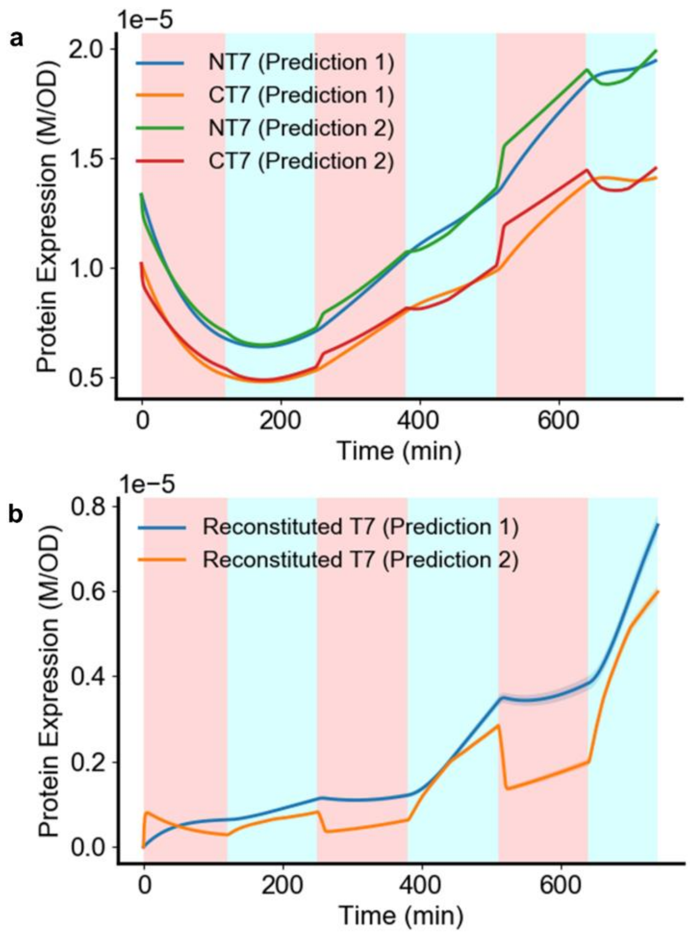


**Supplementary Figure S9.** Model predicted expression profiles for the (a) unbounded NT7 and CT7 protein fragments and (b) reconstituted T7 following the 2hr OFF-ON simulation dynamics, using fitted parameters from continuous data (Prediction 1) and 3hr OFF-ON dynamic data (Prediction 2).

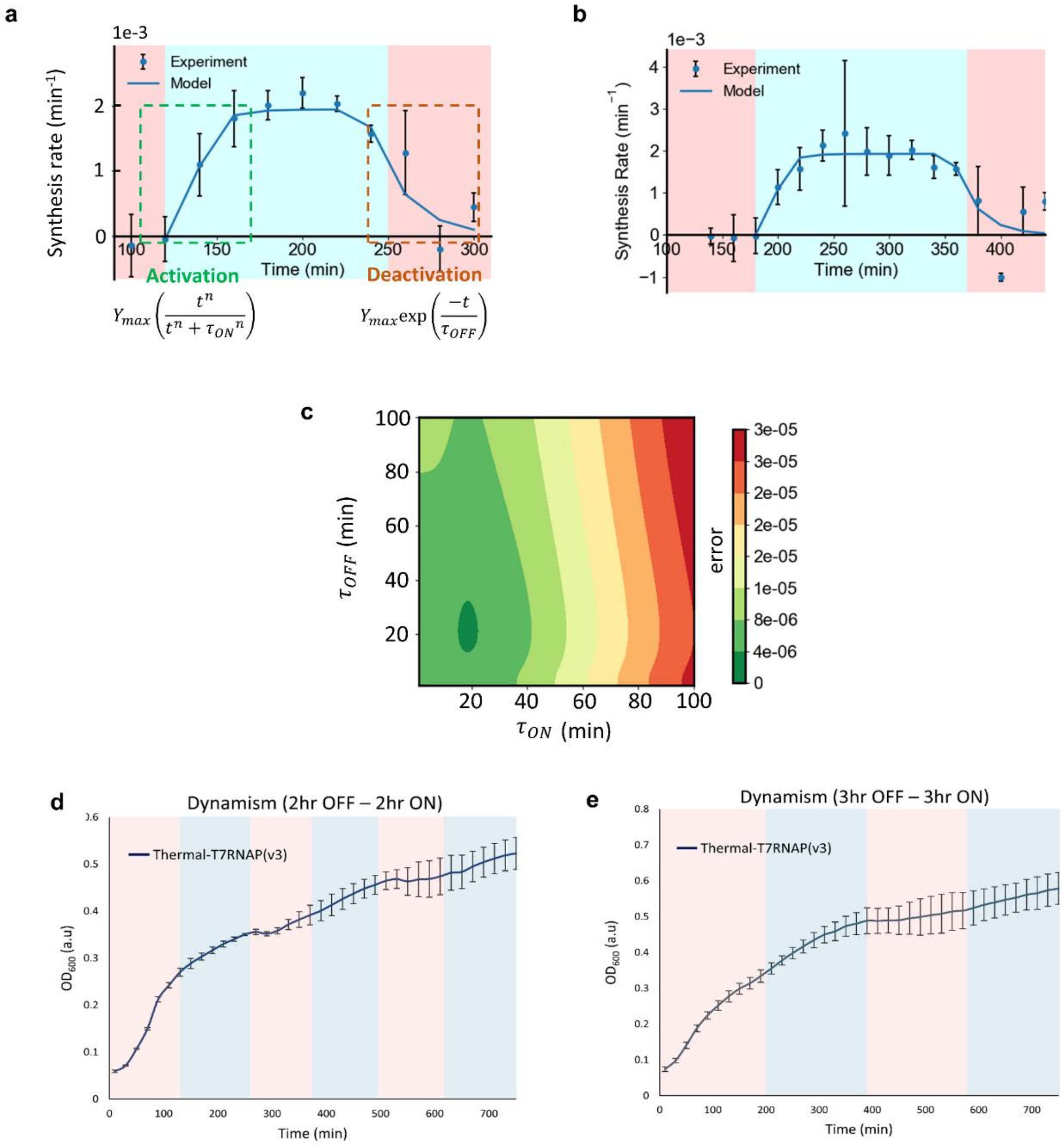


**Supplementary Figure S10.** Derivation of activation and deactivation time constants ($\tau_{ON}$ and $\tau_{OFF}$) of Thermal-T7RNAP(v3) system based on the synthesis rate profiles for the constant (a) 2-hour interval and (b) 3-hour interval thermal cycles, when alternating between 37 °C and 30 °C. The best estimated activation and deactivation time constants ($\tau_{ON}$= 18.82 min, $\tau_{OFF}$= 20.80 min) were derived from the synthesis rate profiles based upon the least-square analysis. In the equations, $\gamma_{max}$ refers to the maximum protein synthesis rate and n denotes the Hill coefficient (n = 4). (c) Heatmap error analysis. Grid search for the narrow range of $\tau_{ON}$ and $\tau_{OFF}$ combination that recapitulates the system transitional kinetics. The heatmap represents the calculated combined sum-squared errors (SSEs) from both the 3-hour and 2-hour interval experimental profiles. The corresponding growth profiles at (d) 2hr interval and (e) 3hr intervals are shown. Data information: All data are represented as mean ± S.D. (n = 3).

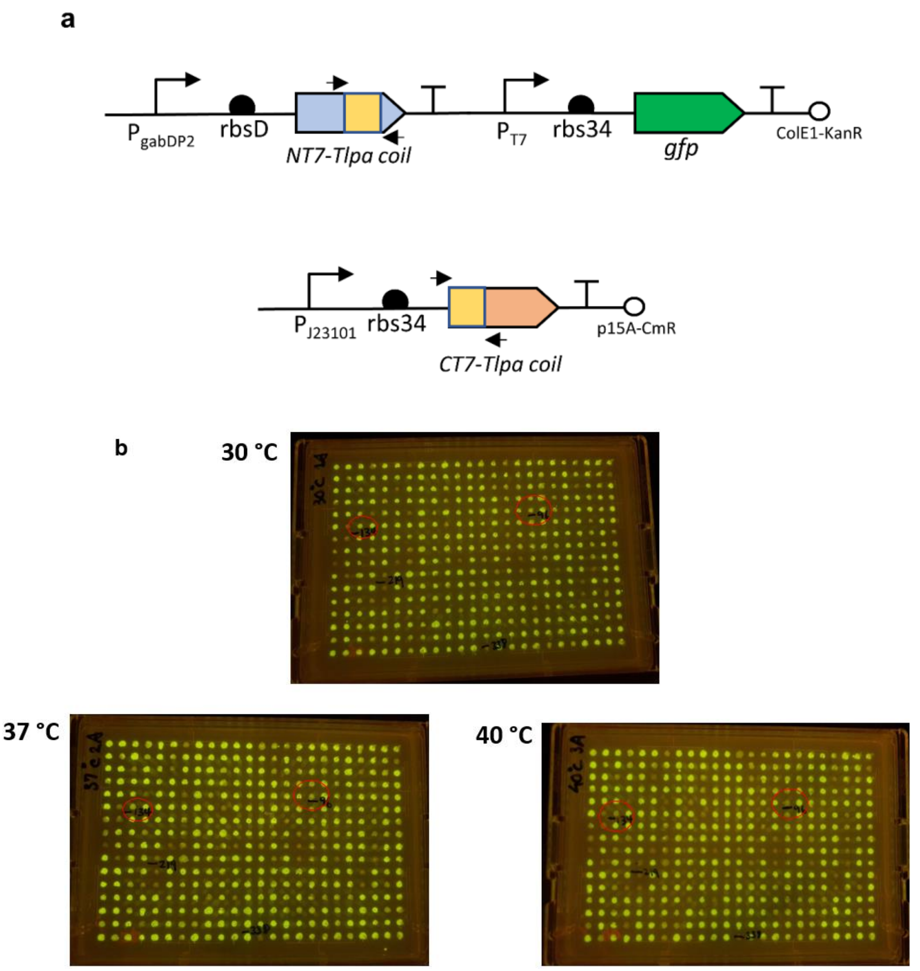


**Supplementary Figure S11.** Mutagenesis framework. (a) Genetic circuits of Thermal-T7RNAP(v1) system where the coiled-coil domains of the NT7 and CT7 protein fragments are mutated (indicated by the primer arrows). (b) Replicated plates after overnight incubation at the respective temperature. The individual colony are each picked and plated by colony picker. The highlighted colonies indicate shortlisted colonies with ideal thermal characteristics ('ON' states at low temperature).

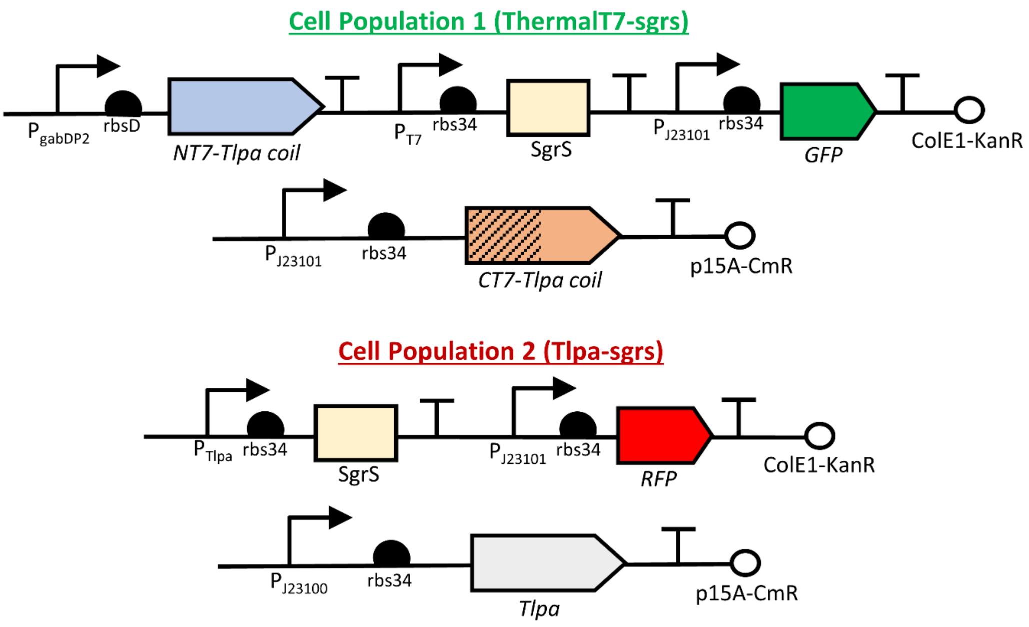


**Supplementary Figure S12.** Genetic circuits for $GFP_{ThermalT7RNAP\text{-}SgrS}$ and $RFP_{Tlpa\text{-}SgrS}$ cells that will make up the co-culture. In the control co-culture, while maintaining similar genetic configuration, the '$p_{T7}$-rbs34-SgrS' or '$p_{Tlpa}$-rbs34-SgrS' cassette are removed.

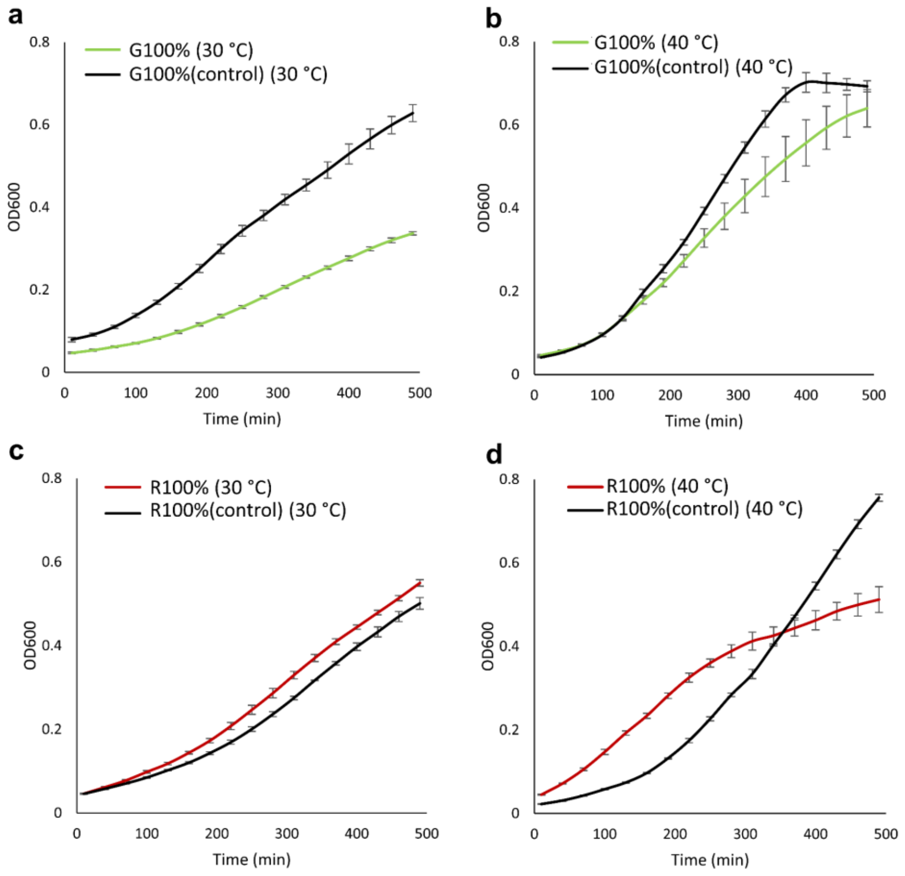


**Supplementary Figure S13.** Continuous growth profiles of the individual monocultures (100%) at 30 °C and 40 °C. (a) $GFP_{ThermalT7RNAP\text{-}SgrS}$ and its (b) GFP control (no SgrS expression) are shown. (c) The $RFP_{Tlpa\text{-}SgrS}$ cells and its (d) RFP control are also shown. Data information: All data are represented as mean ± S.D. (n = 3).

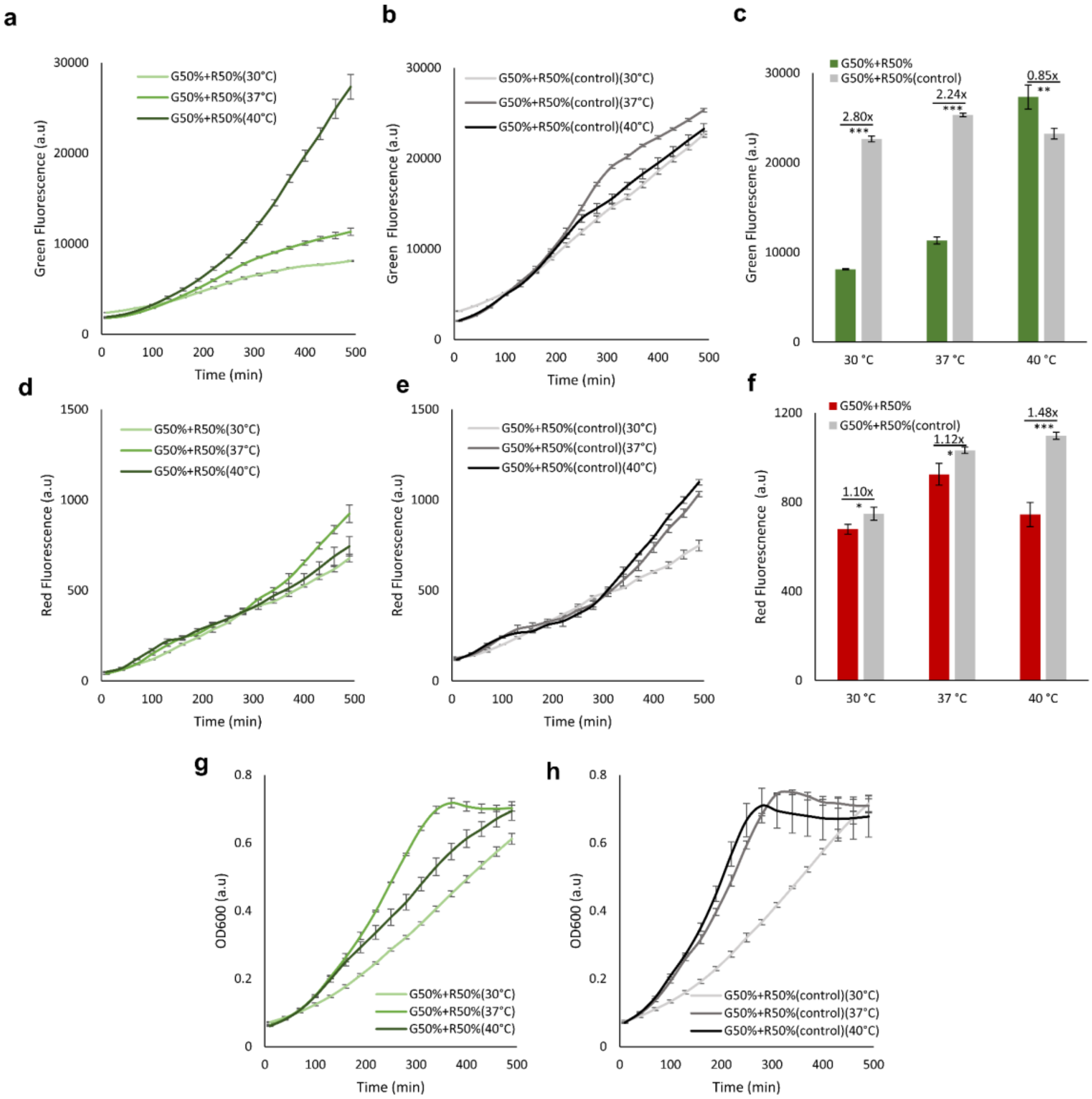


**Supplementary Figure S14.** Characterization of the co-culture at 30 °C, 37 °C and 40 °C. The co-culture harbored the growth inhibition circuitries and seeded at equal initial proportions of GFP and RFP cells (G50%+R50%). The control co-culture (no SgrS expression) was also included. The individual fluorescence for the (a-c) GFP and (d-f) RFP cells were characterized in the microplate reader for 8 hours. Their growth curves were also shown (g-h). Data information: All data are represented as mean ± S.D. (n = 3). Statistical significance of ***P < 0.001, **P<0.01 and *P<0.05 were calculated based on two-sample unpaired t-test.  The corresponding fold-change between the temperatures were shown above each of the bar charts.

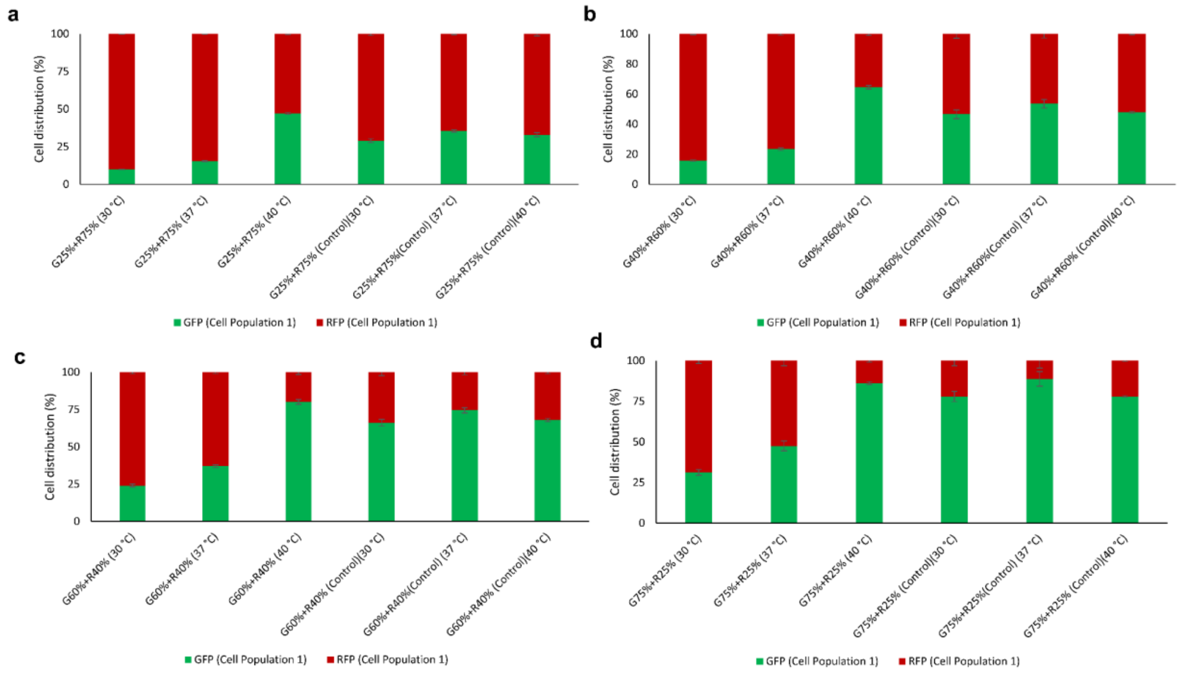


**Supplementary Figure S15.** Final cell distribution of the co-culture after 8hrs. (a-d) The experimental cell distributions for different initial seeding ratios were shown, as derived from the actual fluorescence values. Data information: All data are represented as mean ± S.D. (n = 3).

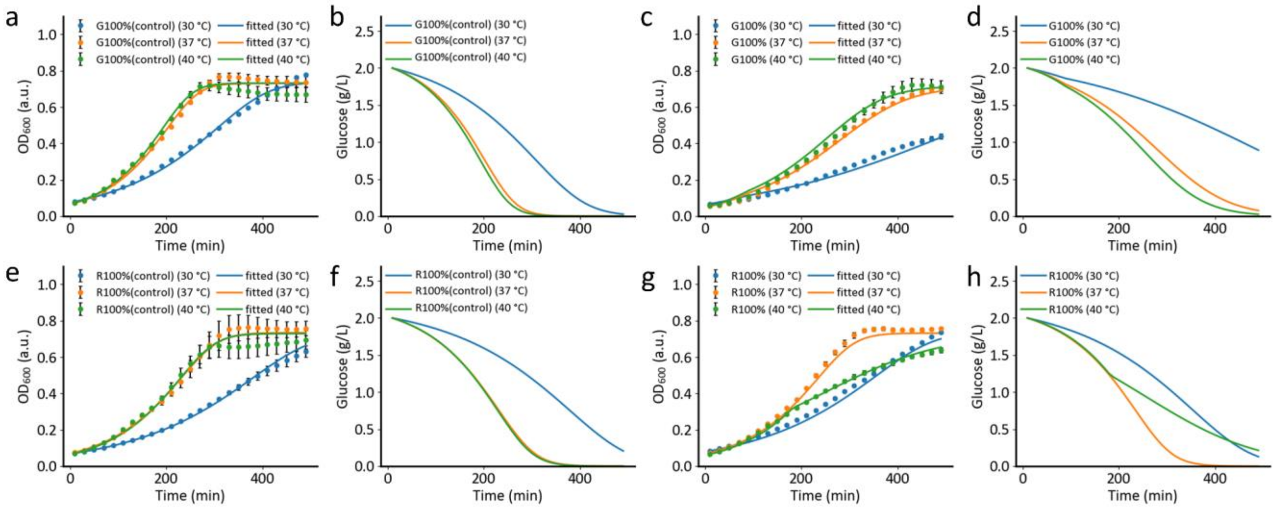


**Supplementary Figure S16.** Model-fitted growth profiles for the monocultures (100%) and their glucose utilization curves at different temperatures (30 °C, 37 °C and 40 °C). (a,b) GFP control cells (no SgrS expression) and (e,f) RFP control cells were shown. And with the introduction of SgrS, (c,d) $GFP_{ThermalT7RNAP\text{-}SgrS\ and}$ and (g,h) $RFP_{Tlpa\text{-}SgrS}$ cells were also shown. The filled circles represent the experimental measurements while the solid lines denote the corresponding model fittings. Data information: All experimental data are represented as mean ± S.D. (n = 3).

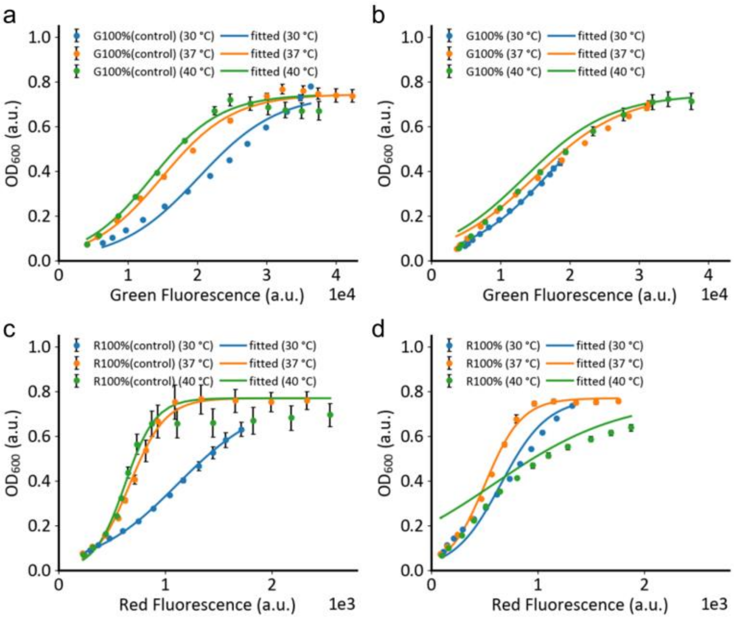


**Supplementary Figure S17.** Correlation between the fluorescence and growth for monocultures (100%) at different temperatures (30 °C, 37 °C and 40 °C) (a) for the GFP control cells; (b) GFP cells containing the growth inhibition module; (c-d) for the RFP control cells and cells with growth inhibition module. The experimental data represented as filled circles were overlay with the model-fitted solid lines. A more phenomenological formulation was used to represent the correlations which could be useful to derive the actual OD values from the measured fluorescence instead of relying on the fluorescence levels alone (Supplementary Table S7-S8).

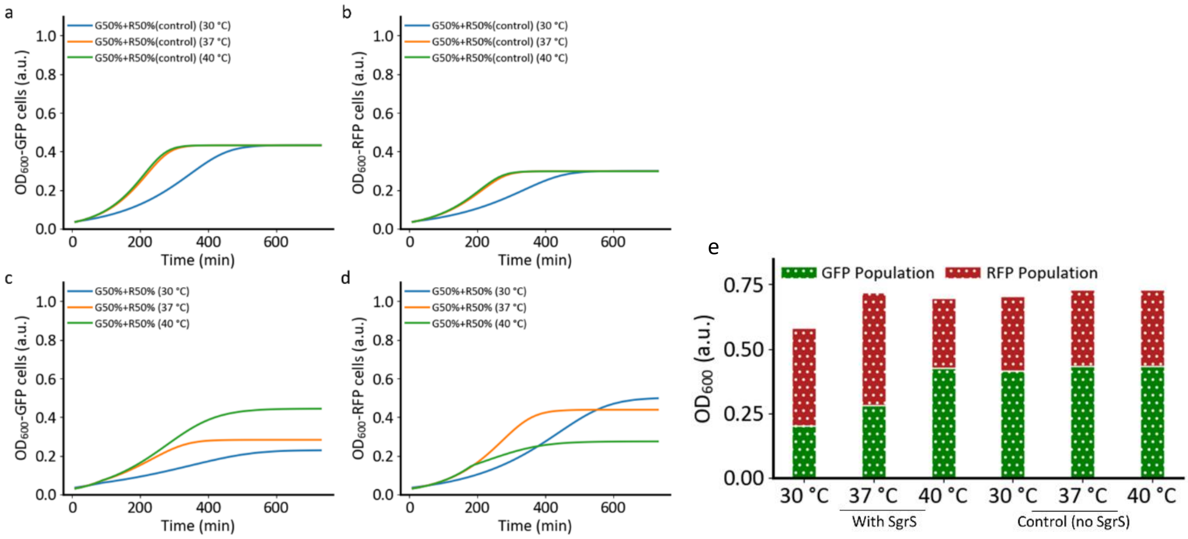


**Supplementary Figure S18.** (a-d) Model predicted individual $OD_{600}$ growth profiles at the three different temperatures (30 °C, 37 °C, 40 °C) for GFP and RFP cell populations within the co-culture at equal initial cell proportions (G50%+R50%) for control and non-control cells. (e) The predicted final OD values for the GFP and RFP cells after 8 hours. Data for both control and non-control cells were shown.

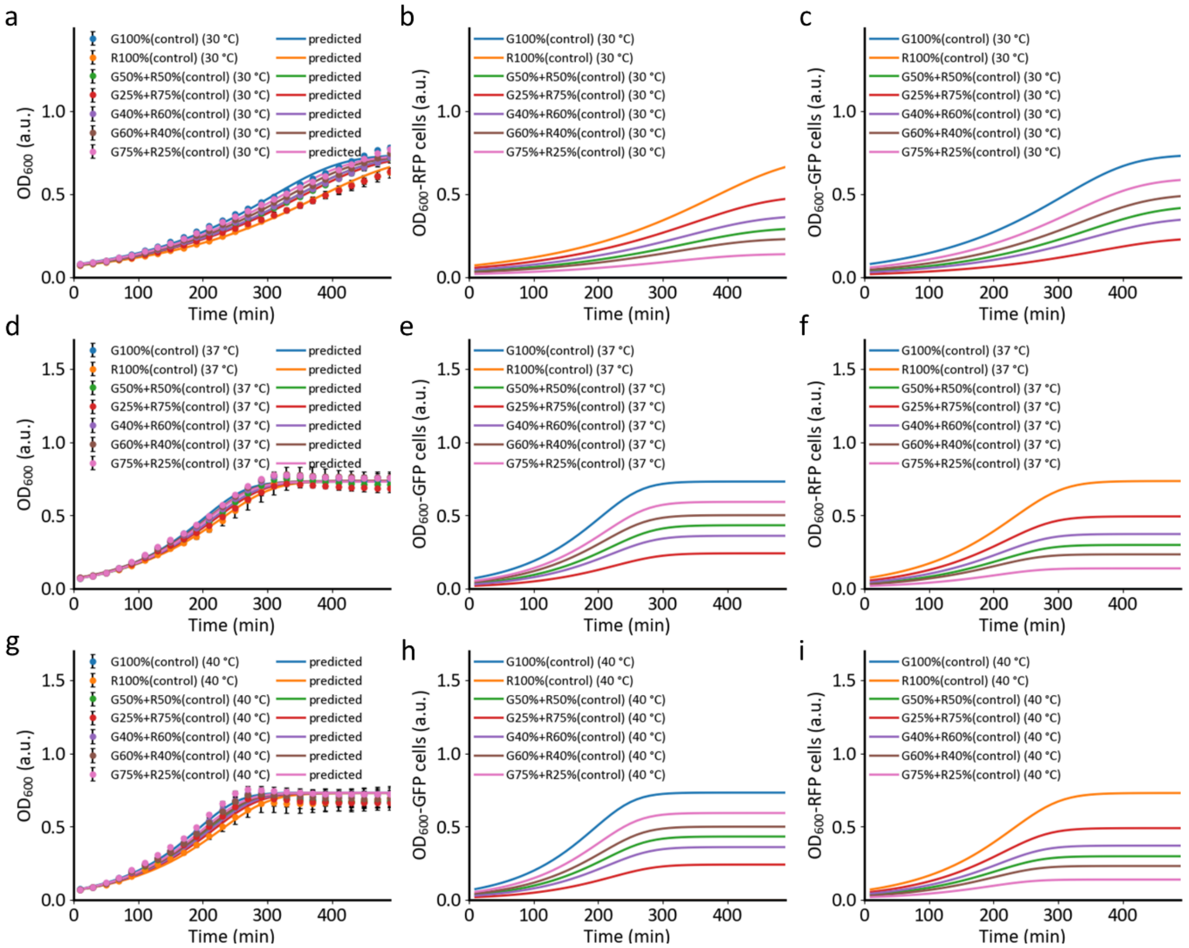


**Supplementary Figure S19.** Growth profiles for control co-cultures when subjected to different initial cell proportions under different temperatures (30 °C, 37 °C, 40 °C). (a, d, g) Combined growth profiles for the co-cultures. (b-c, e-f, h-i) The respective model-predicted cell growth profiles for the individual cell populations within the co-cultures. The filled circles represent the experimental measurements (N=3) while solid lines denote the corresponding model predictions.

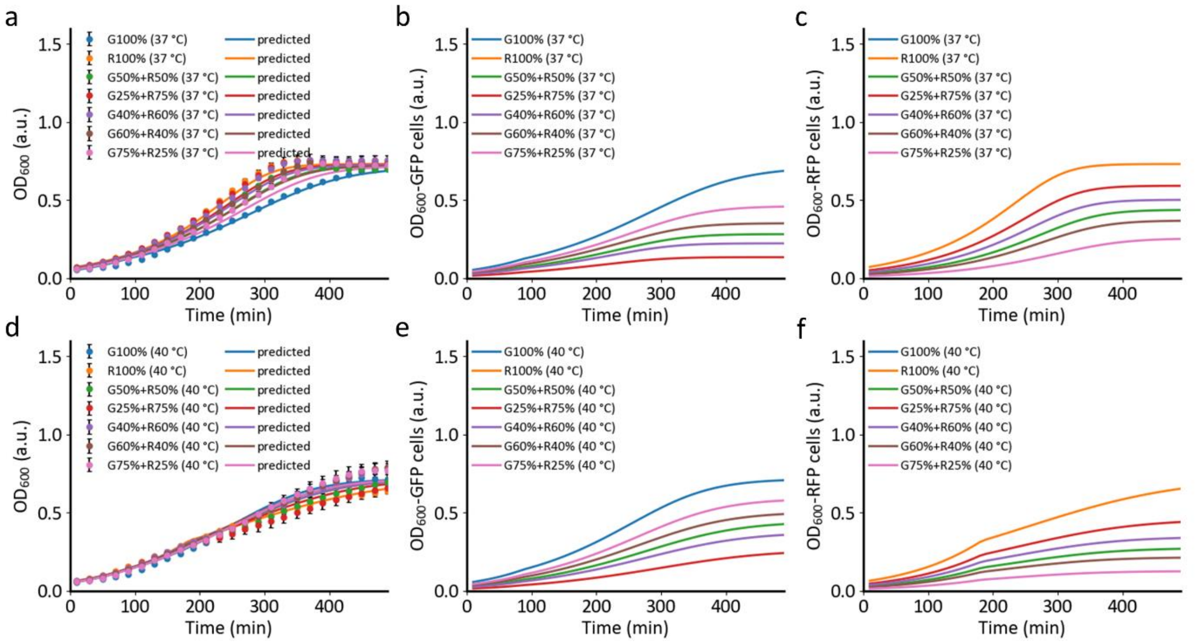


**Supplementary Figure S20.** Growth profiles for the co-cultures with growth inhibition modules when subjected to different initial cell proportions under different temperatures (37 °C and 40 °C). (a, d) Combined growth profiles (b-c, e-f) The respective model-predicted growth profiles for the individual cell populations within the co-cultures. The filled circles represent the experimental measurements (N=3) while solid lines denote the corresponding model predictions.

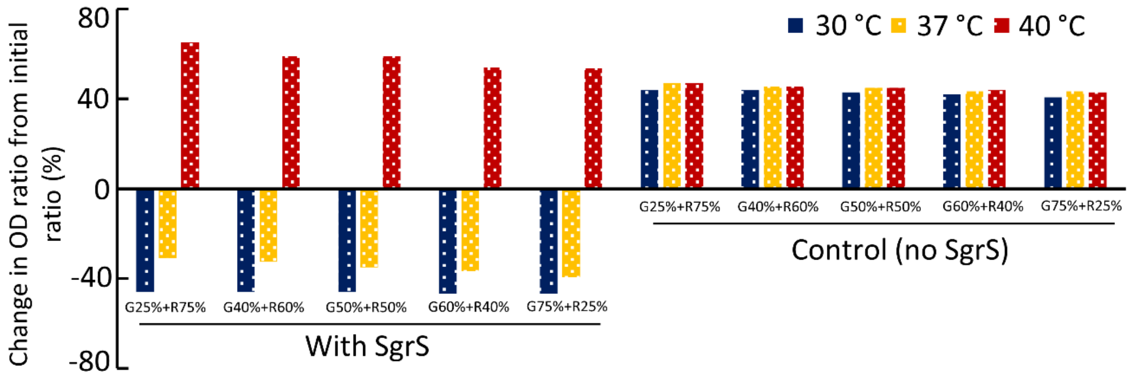


**Supplementary Figure S21.** Percentage change in final predicted OD ratios (after 8 hrs) from the various initial ratios for the co-cultures (with growth inhibition module) and control co-cultures (no SgrS expression).

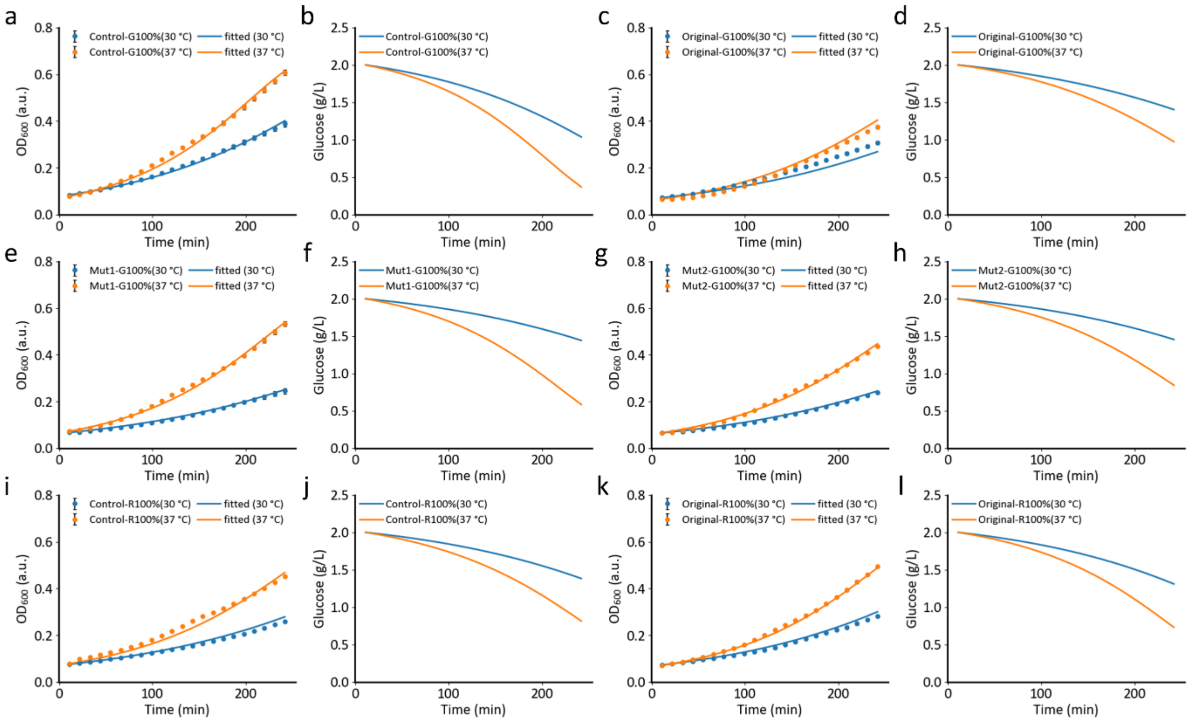


**Supplementary Figure S22.** In monoculture condition (100%), individual growth profiles and glucose utilization curves for the (a-b) GFP control and (i-j) RFP control cells were shown. In the cells with growth inhibition module, the (c-d, k-l) wild-type ThermalT7-SgrS (original) and the mutant systems (e-f) Mut 1 and (g-h) Mut 2 were also shown. The filled circles represent the experimental measurements (N=3) while solid lines denote the corresponding model fittings.

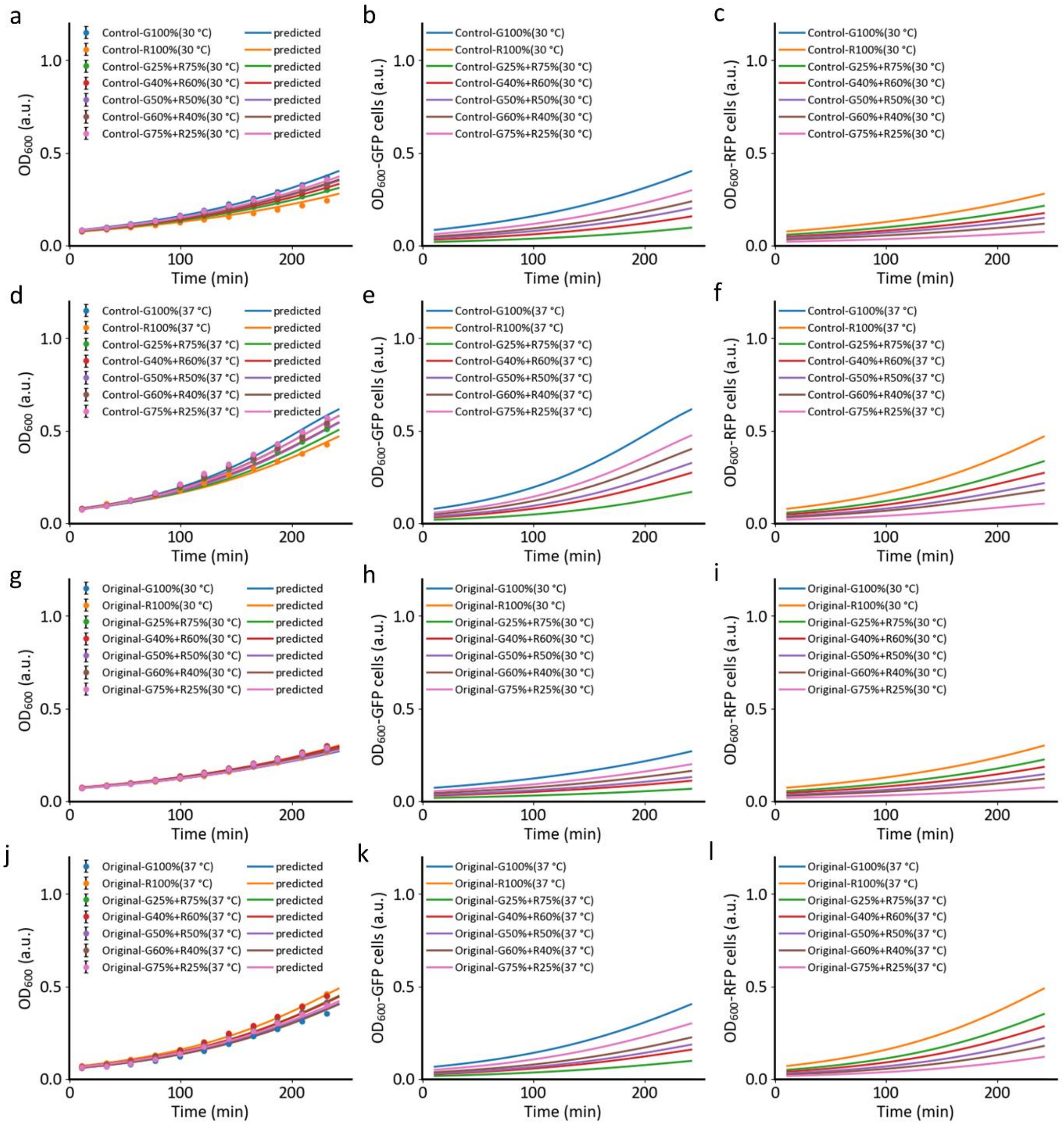


**Supplementary Figure S23.** Combined growth profiles for the (a, d) control co-culture (no SgrS expression) and (g, j) co-culture (with growth inhibition modules) with different initial cell proportions under 30 °C and 37 °C. The respective model-predicted cell growth profiles for the individual cell populations within the (b-f) control co-culture and (h-l) co-cultures (with SgrS expression) were also shown. The filled circles represent the experimental measurements (N=3) while solid lines denote the corresponding model predictions.

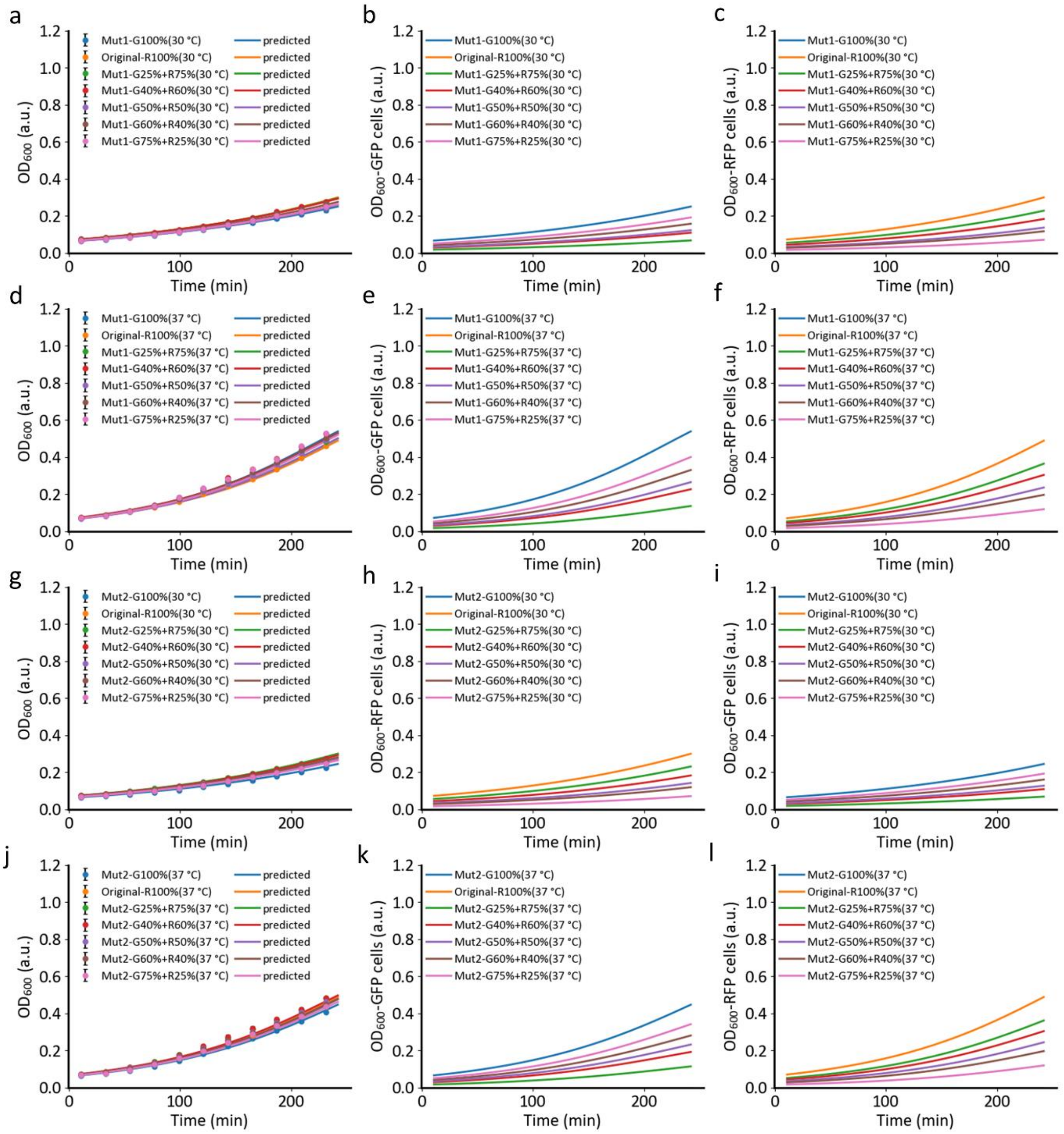


**Supplementary Figure S24.** Combined growth profiles for the (a, d) mutant co-culture 1 (Mut1) and (g, j) mutant co-culture 2 (Mut 2) with different initial cell proportions under 30 °C and 37 °C. The respective model-predicted cell growth profiles for the individual cell populations within the (b-f) Mut 1 co-culture and (h-l) Mut 2 co-cultures were also shown. The filled circles represent the experimental measurements (N=3) while solid lines denote the corresponding model predictions.

### SUPPLEMENTARY NOTES

**Supplementary Note S1:**

We observed different correlation profiles between the reporting fluorescence expression and the cell growth at different temperatures in the various monocultures (Supplementary Figure S17). Both GFP and RFP control cells exhibited trends of decreasing fluorescence with increasing temperatures, as supported by an increase in their half-activation coefficients and curve gradients (Supplementary Table S7-S8). When equipped with the growth inhibition systems, the correlations were also found to be disparate for both $GFP_{ThermalT7RNAP\text{-}SgrS}$ and $RFP_{Tlpa\text{-}SgrS}$ cells as compared to their respective controls. All the half-activation coefficients and the curve gradients at different temperatures have been shifted (Supplementary Table S8). This shows that the thermal growth modulation systems which express the growth inhibitory SgrS at different temperatures may have shifted the resource allocation between the cell OD growth and the fluorescence protein expression.

**Supplementary Note S2:**

Bayesian parameter inference method was used to infer the probability distributions of some parameter estimates. Based on the Bayes' theorem, the posterior is expressed as follows:

$$P(\theta|D) = \frac{P(D|\theta).P(\theta)}{P(D)} \quad \text{(Eq. S29)}$$

$P(\theta|D)$ denotes the joint conditional probability of different parameter combinations given the experimental measured data. The $P(D|\theta)$ computes the likelihood, $P(\theta)$ represents the prior regardless of the measured data and $P(D)$ is the evidence or data regardless of the event. The general acceptance rule is governed by

$$\sum\nolimits_{i}^{n} -Log\left(\sigma\sqrt{2\pi}\right) - \frac{SSE}{2\sigma^2} + Log\left(prior(\mu_{new}, \sigma_{new})\right) > \sum\nolimits_{i}^{n} -Log\left(\sigma\sqrt{2\pi}\right) - \frac{SSE}{2\sigma^2} + Log\left(prior(\mu_{current}, \sigma_{current})\right) \quad \text{(Eq. S30)}$$

The logarithmic form is used to ensure numerical stability. The $SSE$ represents the sum of squared errors between the experimental observations and model simulations, and $\sigma$ is chosen to be 1e-6 to ensure proper convergence time. If the above condition is not satisfied or the ratio is not larger than 1, the ratio is then compared to a uniformly generated random number within [0, 1]. The new set of parameters $\theta_{new}$ will be accepted if the ratio is larger than the random number, otherwise the $\theta_{new}$ will be rejected, and the $\theta_{current}$ will be kept. The number of iterations and the transition model were chosen carefully to ensure a proper convergence has been reached within a short computation time. The mean values of the priors were used as the starting sample.

### SUPPLEMENTARY REFERENCE

1. Liao, C., Blanchard, A.E. and Lu, T. (2017) An integrative circuit–host modelling framework for predicting synthetic gene network behaviours. *Nature Microbiology*, **2**, 1658-1666.
